# Quantifying within-host diversity of H5N1 influenza viruses in humans and poultry in Cambodia

**DOI:** 10.1101/683151

**Authors:** Louise H. Moncla, Trevor Bedford, Philippe Dussart, Srey Viseth Horm, Sareth Rith, Philippe Buchy, Erik A Karlsson, Lifeng Li, Yongmei Liu, Huachen Zhu, Yi Guan, Thomas C. Friedrich, Paul F. Horwood

**Affiliations:** Fred Hutchinson Cancer Research Center, Seattle, Washington, United States; University of Washington, Seattle, Washington, United States; Virology Unit, Institut Pasteur du Cambodge, Institut Pasteur International Network, Phnom Penh, Cambodia; GlaxoSmithKline, Vaccines R&D, Singapore, Singapore; Joint Influenza Research Centre (SUMC/HKU), Shantou University Medical College, Shantou, People’s Republic of China; State Key Laboratory of Emerging Infectious Diseases/Centre of Influenza Research, School of Public Health, The University of Hong Kong, Hong Kong, SAR, People’s Republic of China; Department of Pathobiological Sciences, University of Wisconsin School of Veterinary Medicine, Madison, WI, United States; Wisconsin National Primate Research Center, Madison, WI, United States; College of Public Health, Medical and Veterinary Sciences, James Cook University, Townsville, Australia

## Abstract

Avian influenza viruses (AIVs) periodically cross species barriers and infect humans. The likelihood that an AIV will evolve mammalian transmissibility depends on acquiring and selecting mutations during spillover, but data from natural infection is limited. We analyze deep sequencing data from infected humans and domestic ducks in Cambodia to examine how H5N1 viruses evolve during spillover. Overall, viral populations in both species are predominated by low-frequency (<10%) variation shaped by purifying selection and genetic drift, and half of the variants detected within-host are never detected on the H5N1 virus phylogeny. However, we do detect a subset of mutations linked to human receptor binding and replication (PB2 E627K, HA A150V, and HA Q238L) that arose in multiple, independent humans. PB2 E627K and HA A150V were also enriched along phylogenetic branches leading to human infections, suggesting that they are likely human-adaptive. Our data show that H5N1 viruses generate putative human-adapting mutations during natural spillover infection, many of which are detected at >5% frequency within-host. However, short infection times, genetic drift, and purifying selection likely restrict their ability to evolve extensively during a single infection. Applying evolutionary methods to sequence data, we reveal a detailed view of H5N1 virus adaptive potential, and develop a foundation for studying host-adaptation in other zoonotic viruses.

**Author summary:** H5N1 avian influenza viruses can cross species barriers and cause severe disease in humans. H5N1 viruses currently cannot replicate and transmit efficiently among humans, but animal infection studies and modeling experiments have suggested that human adaptation may require only a few mutations. However, data from natural spillover infection has been limited, posing a challenge for risk assessment. Here, we analyze a unique dataset of deep sequence data from H5N1 virus-infected humans and domestic ducks in Cambodia. We find that well-known markers of human receptor binding and replication arise in multiple, independent humans. We also find that 3 mutations detected within-host are enriched along phylogenetic branches leading to human infections, suggesting that they are likely human-adapting. However, we also show that within-host evolution in both humans and ducks are shaped heavily by purifying selection and genetic drift, and that a large fraction of within-host variation is never detected on the H5N1 phylogeny. Taken together, our data show that H5N1 viruses do generate human-adapting mutations during natural infection. However, short infection times, purifying selection, and genetic drift may severely limit how much H5N1 viruses can evolve during the course of a single infection.

## Introduction

Influenza virus cross-species transmission poses a continual threat to human health. Since emerging in 1997, H5N1 avian influenza viruses (AIVs) have caused 860 confirmed infections and 454 deaths in humans[1]. H5N1 viruses naturally circulate in aquatic birds, but some lineages have integrated into poultry populations. H5N1 viruses are now endemic in domestic birds in some countries[2–4], and concern remains that continued human infection may one day facilitate human adaptation.

The likelihood that an AIV will adapt to replicate and transmit among humans depends on generating and selecting human-adaptive mutations during spillover. Influenza viruses have high mutation rates[5–8], short generation times[9], and large populations, and rapidly generate diversity within-host. Laboratory studies using animal models[10–12] show that only 3-5 amino acid substitutions may be required to render H5N1 viruses mammalian-transmissible[10–12], and that viral variants present at frequencies as low as 5% may be transmitted by respiratory droplets[13]. Subsequent modeling studies suggest that within-host dynamics are conducive to generating human-transmissible viruses, but that these viruses may remain at frequencies too low for transmission[14, 15]. Although these studies offer critical insight for H5N1 virus risk assessment, it is unclear whether they adequately describe how cross-species transmission proceeds in nature.

H5N1 virus outbreaks offer rare opportunities to study natural cross-species transmission, but data are limited. One study of H5N1 virus-infected humans in Vietnam identified mutations affecting receptor binding, polymerase activity, and interferon antagonism; however, they remained at low frequencies throughout infection[16]. Recent characterization of H5N1 virus-infected humans in Indonesia identified novel mutations within-host that enhance polymerase activity in human cells[17]. Unfortunately, neither of these studies include data from naturally infected poultry, which would provide a critical comparison for assessing whether infected humans exhibit signs of adaptive evolution. A small number of studies have examined within-host diversity in experimentally infected poultry[18–20], but these may not recapitulate the dynamics of natural infection.

As part of ongoing diagnostic and surveillance effort, the Institut Pasteur du Cambodge collects and confirms samples from AIV-infected poultry during routine market surveillance, and from human cases and poultry during AIV outbreaks. Since H5N1 was first detected in Cambodia in 2004, 56 human cases and 58 poultry outbreaks have been confirmed and many more have gone undetected[21]. Here we analyze previously generated deep sequence data[22] from 8 infected humans and 5 infected domestic ducks collected in Cambodia between 2010 and 2014. We find that viral populations in both species are dominated by low-frequency variation shaped by population expansion, purifying selection, and genetic drift. We identify a handful of mutations in humans linked to improved mammalian replication and transmissibility, two of which were detected in multiple samples, suggesting that adaptive mutations arise during natural spillover infection. Although most within-host mutations are not linked to human infections on the H5N1 virus phylogeny, three mutations identified within-host are enriched on phylogenetic branches leading to human infections. Our data suggest that known adaptive mutations do occur in natural H5N1 virus infection, but that a short duration of infection, randomness, and purifying selection may together limit the evolutionary capacity of these viruses to evolve extensively during any individual spillover event.

## Methods

### Viral sample collection

The Institute Pasteur in Cambodia is a World Health Organization H5 Reference Laboratory (H5RL) and has a mandate to assist the Cambodian Ministry of Health and the Ministry of Agriculture, Forestry, and Fisheries in conducting investigations into human cases and poultry outbreaks of H5N1 virus, respectively. Surveillance for human cases of H5N1 virus infection is conducted through influenza-like-illness, severe acute respiratory illness, and event-based surveillance in a network of hospitals throughout the country [23]. Poultry outbreaks of H5N1 virus are detected through passive surveillance following reports from farmers and villagers of livestock illness or death. The H5RL conducts confirmation of H5N1 virus detection and further characterization (genetic and antigenic) of H5N1 virus strains.

### Human subjects and IRB approval

The Cambodian influenza surveillance system is a public health activity managed by the Ministry of Health in Cambodia and has a standing authorization from the National Ethics Committee for Human Research. The deep sequence analysis of H5N1 influenza virus from human samples was approved for this study by the National Ethics Committee for Human Research (#266NECHR).

### RNA isolation and RT-qPCR

RNA was extracted from swab samples using the QIAmp Viral RNA Mini Kit (Qiagen, Valencia, CA, USA), following manufacturer’s guidelines and eluted in buffer AVE. Extracts were tested for influenza A virus (M-gene)[24] and subtypes H5 (primer sets H5a and H5b), N1, H7, and H9 by using quantitative RT-PCR (qRT-PCR) using assays sourced from the International Reagent Resource (https://www.internationalreagentresource.org/Home.aspx), as previously outlined[25]. Only samples with high viral load (≥10^3^ copies/μl of extracted viral RNA in buffer AVE), as assessed by RT-qPCR, were selected for sequence analysis. All samples were sequenced directly from the original specimen, without passaging in cell culture or eggs. Information on the samples included in the present analyses are presented in Table 1. **cDNA generation and PCR** cDNA was generated using Superscript IV Reverse Transcriptase (Invitrogen, Carlsbad, CA, USA) and custom influenza primers targeting the conserved ends for whole genome amplification[26]. The following primers were pooled together in a 1.5 : 0.5 : 2.0 : 1.0 ratio: Uni-1.5: ACGCGTGATCAGCAAAAGCAGG, Uni-0.5: ACGCGTGATCAGCGAAAGCAGG, Uni-2.0: ACGCGTGATCAGTAGAAACAAGG, and Uni-1.0: AGCAAAAGCAGG. 1 μl of this primer pool was added to 1 μl of 10 mM dNTP mix (Invitrogen) and 11 μl of RNA. Contents were briefly mixed and heated for 5 minutes at 65°C, followed by immediate incubation on ice for at least 1 minute. Next, a second mastermix was made with 4 μl of 5X Superscript IV Buffer, 1 μl of 100 mM DTT, 1 μl of RNaseOut Recombinant RNase Inhibitor, and 1 μl of SuperScript IV Reverse Transcriptase (200 U/μl) (Invitrogen). 7 μl of mastermix was added to each sample, for a total volume of 20 μl. This mixture was briefly mixed, incubated at 55°C for 20 minutes, then inactivated by incubating at 80°C for 10 minutes. Whole genomic amplification of the influenza virus was conducted using Ex Taq™ Hot Start Version (TaKaRa). Forward primers were Uni-1.5 and Uni-0.5 mixed in a ratio of 3:2, and reverse primer was Uni-2.0. The temperature cycle parameters were 98°C for 2 min, and then 5 cycles (98°C for 30 seconds, 45°C for 30 seconds, and 72°C for 3 minutes), followed by 25 cycles (98°C for 30 seconds, 55°C for 30 seconds, and 72°C for 3 minutes).

**Table 1:** Sample information.

| Sample ID | Host | Sample type | Collection | Date | Days post-symptom onset | vRNA copies/μl (after vRNA extraction) | Clade |
| --- | --- | --- | --- | --- | --- | --- | --- |
| A/duck/Cambodia/PV027D1/2010 | Domestic duck | Pooled organs | Poultry outbreak investigation | April 2010 | NA | $5.45 \times 10^6$ | 1.1.2 |
| A/duck/Cambodia/083D1/2011 | Domestic duck | Pooled organs | Poultry outbreak investigation | September 2011 | NA | $3.74 \times 10^7$ | 1.1.2 |
| A/duck/Cambodia/381W11M4/2013 | Domestic duck | Pooled throat and cloacal swab | Live bird market surveillance | March 2013 | NA | $7.37 \times 10^5$ | 1.1.2/2.3.2.1a reassortant |
| A/duck/Cambodia/Y0224301/2014 | Domestic duck | Pooled organs | Poultry outbreak investigation | February 2014 | NA | $2.0 \times 10^5$ | 1.1.2/2.3.2.1a reassortant |
| A/duck/Cambodia/Y0224304/2014 | Domestic duck | Pooled organs | Poultry outbreak investigation | February 2014 | NA | $5.0 \times 10^6$ | 1.1.2/2.3.2.1a reassortant |
| A/Cambodia/V0401301/2011 | Human (10F, died) | Throat swab | Event-based surveillance | April 2011 | 9 | $5.02 \times 10^3$ | 1.1.2 |
| A/Cambodia/V0417301/2011 | Human<br>(5F, died) | Throat<br>swab | Event-based<br>surveillance | April 2011 | 5 | $8.98 \times 10^4$ | 1.1.2 |
| A/Cambodia/W0112303/2012 | Human<br>(2M, died) | Throat<br>swab | Event-based<br>surveillance | January<br>2012 | 7 | $2.05 \times 10^3$ | 1.1.2 |
| A/Cambodia/X0125302/2013 | Human<br>(1F, died) | Throat<br>swab | Event-based<br>surveillance | January<br>2013 | 12 | $6.84 \times 10^4$ | 1.1.2/2.3.2.1a<br>reassortant |
| A/Cambodia/X0128304/2013 | Human<br>(9F, died) | Throat<br>swab | Event-based<br>surveillance | January<br>2013 | 8 | $5.09 \times 10^3$ | 1.1.2/2.3.2.1a<br>reassortant |
| A/Cambodia/X0207301/2013 | Human<br>(5F, died) | Throat<br>swab | Event-based<br>surveillance | February<br>2013 | 12 | $1.73 \times 10^5$ | 1.1.2/2.3.2.1a<br>reassortant |
| A/Cambodia/X0219301/2013 | Human<br>(2M, died) | Throat<br>swab | Event-based<br>surveillance | February<br>2013 | 12 | $1.66 \times 10^3$ | 1.1.2/2.3.2.1a<br>reassortant |
| A/Cambodia/X1030304/2013 | Human<br>(2F, died) | Throat<br>swab | Event-based<br>surveillance | October<br>2013 | 8 | $1.08 \times 10^4$ | 1.1.2/2.3.2.1a<br>reassortant |

### Library preparation and sequencing

For each sample, amplicons were quantified using the Qubit^TM^ dsDNA BR Assay Kit (Invitrogen), pooled in equimolar concentrations, and fragmented using the NEBNext dsDNA Fragmentase (New England BioLabs, Ipswich, MA). DNA fragments with the size of 350-700 bp were separated on an agarose gel during electrophoresis and purified for input into the NEBNext Ultra DNA Library Prep Kit for Illumina® (New England BioLabs). Prepared libraries were quantified using KAPA Library Quantification Kits for Illumina® platforms (KAPA Biosystems) and pooled in equimolar concentrations to a final concentration of 4 nM, and run using an MiSeq Reagent Kit v2 (Illumina, San Diego, CA) for 500 cycles (2 x 250 bp). Demultiplexed files were output in FASTQ format.

### Processing of raw sequence data, mapping, and variant calling

Human reads were removed from raw FASTQ files by mapping to the human reference genome GRCH38 with bowtie2[27] version 2.3.2 (http://bowtie-bio.sourceforge.net/bowtie2/index.shtml). Reads that did not map to human genome were output to separate FASTQ files and used for all subsequent analyses. Illumina data was analyzed using the pipeline described in detail at https://github.com/lmoncla/illumina_pipeline. Briefly, raw FASTQ files were trimmed using Trimmomatic[28] (http://www.usadellab.org/cms/?page=trimmomatic), trimming in sliding windows of 5 base pairs and requiring a minimum Q-score of 30. Reads that were trimmed to a length of <100 base pairs were discarded. Trimming was performed with the following command: java -jar Trimmomatic-0.36/trimmomatic-0.36.jar SE input.fastq output.fastq SLIDINGWINDOW:5:30 MINLEN:100. Trimmed reads were mapped to consensus sequences previously derived[22] using bowtie2[27] version 2.3.2 (http://bowtie-bio.sourceforge.net/bowtie2/index.shtml), using the following command: bowtie2 -x reference_sequence.fasta -U read1.trimmed.fastq,read2.trimmed.fastq -S output.sam --local. Duplicate reads were removed with Picard (http://broadinstitute.github.io/picard/) with: java -jar picard.jar MarkDuplicates I=input.sam O=output.sam REMOVE_DUPLICATES=true. Mapped reads were imported into Geneious (https://www.geneious.com/) for visual inspection and consensus calling. Consensus sequences were called by reporting the majority base at each site. For nucleotide sites with <100x coverage, a consensus base was not reported, and was instead reported as an “N”. To avoid issues with mapping to an improper reference sequence, we then remapped each sample’s trimmed FASTQ files to its own consensus sequence. These bam files were again manually inspected in Geneious, and a final consensus sequence was called. We were able to generate full-genome data for all samples except for A/Cambodia/X0128304/2013, for which we were lacked data for PB1. These BAM files were then exported and converted to mpileup files with samtools[29] (http://samtools.sourceforge.net/), and within-host variants were called using VarScan[30, 31] (http://varscan.sourceforge.net/). For a variant to be reported, we required the variant site to be sequenced to a depth of at least 100x with a minimum, mean PHRED quality score of 30, and for the variant to be detected in both forward and reverse reads at a frequency of at least 1%. We called variants using the following command: java -jar VarScan.v2.3.9.jar mpileup2snp input.pileup --min-coverage 100 --min-avg-qual 30 --min-var-freq 0.01 --strand-filter 1 --output-vcf 1 > output.vcf. VCF files were parsed and annotated with coding region changes using custom software available here (https://github.com/blab/h5n1-cambodia/blob/master/scripts/H5N1_vcf_parser.py). All amino acid changes for HA are reported and plotted using native H5 numbering, including the signal peptide, which is 16 amino acids in length. For ease of comparison, some amino acid changes are also reported with mature H5 peptide numbering in the manuscript when indicated.

### Phylogenetic reconstruction

We downloaded all currently available H5N1 virus genomes from the EpiFlu Database of the Global Initiative for Sharing All Influenza Data[32, 33] (GISAID, https://www.gisaid.org/) and all currently available full H5N1 virus genomes from the Influenza Research Database (IRD, http://www.fludb.org)[34] and added consensus genomes from our 5 duck samples and 8 human samples. Sequences and metadata were cleaned and organized using fauna (https://github.com/nextstrain/fauna), a database system part of the Nextstrain platform. Sequences were then processed using Nextstrain’s augur software[35] (https://github.com/nextstrain/augur). Sequences were filtered by length to remove short sequences using the following length filters: PB2: 2100 bp, PB1: 2100 bp, PA: 2000 bp, HA: 1600 bp, NP: 1400 bp, NA: 1270 bp, MP: 900 bp, and NS: 800 bp. We excluded sequences with sample collection dates prior to 1996, and those for which the host was annotated as laboratory derived, ferret, or unknown. We also excluded sequences for which the country or geographic region was unknown. Sequences for each gene were aligned using MAFFT[36], and then trimmed to the reference sequence. We chose the A/Goose/Guangdong/1/96(H5N1) genome (GenBank accession numbers: AF144300-AF144307) as the reference genome. IQTREE[37, 38] was then used to infer a maximum likelihood phylogeny, and TreeTime[39] was used to infer a molecular clock and temporally-resolved phylogeny. Tips which fell outside of 4 standard deviations away from the inferred molecular clock were removed. Finally, TreeTime[39] was used to infer ancestral sequence states at internal nodes and the geographic migration history across the phylogeny. We inferred migration among 9 defined geographic regions, China, Southeast Asia, South Asia, Japan and Korea, West Asia, Africa, Europe, South America, and North America, as shown by color in Fig. 1 and **Fig. S2.** Our final trees are available at https://github.com/blab/h5n1-cambodia/tree/master/data/tree-jsons, and include the following number of sequences: PB2: 4063, PB1: 3867, PA: 4082, HA: 6431, NP: 4070, NA: 5357, MP: 3940, NS: 3678. Plotting was performed using baltic (https://github.com/evogytis/baltic).

**Figure 1:**
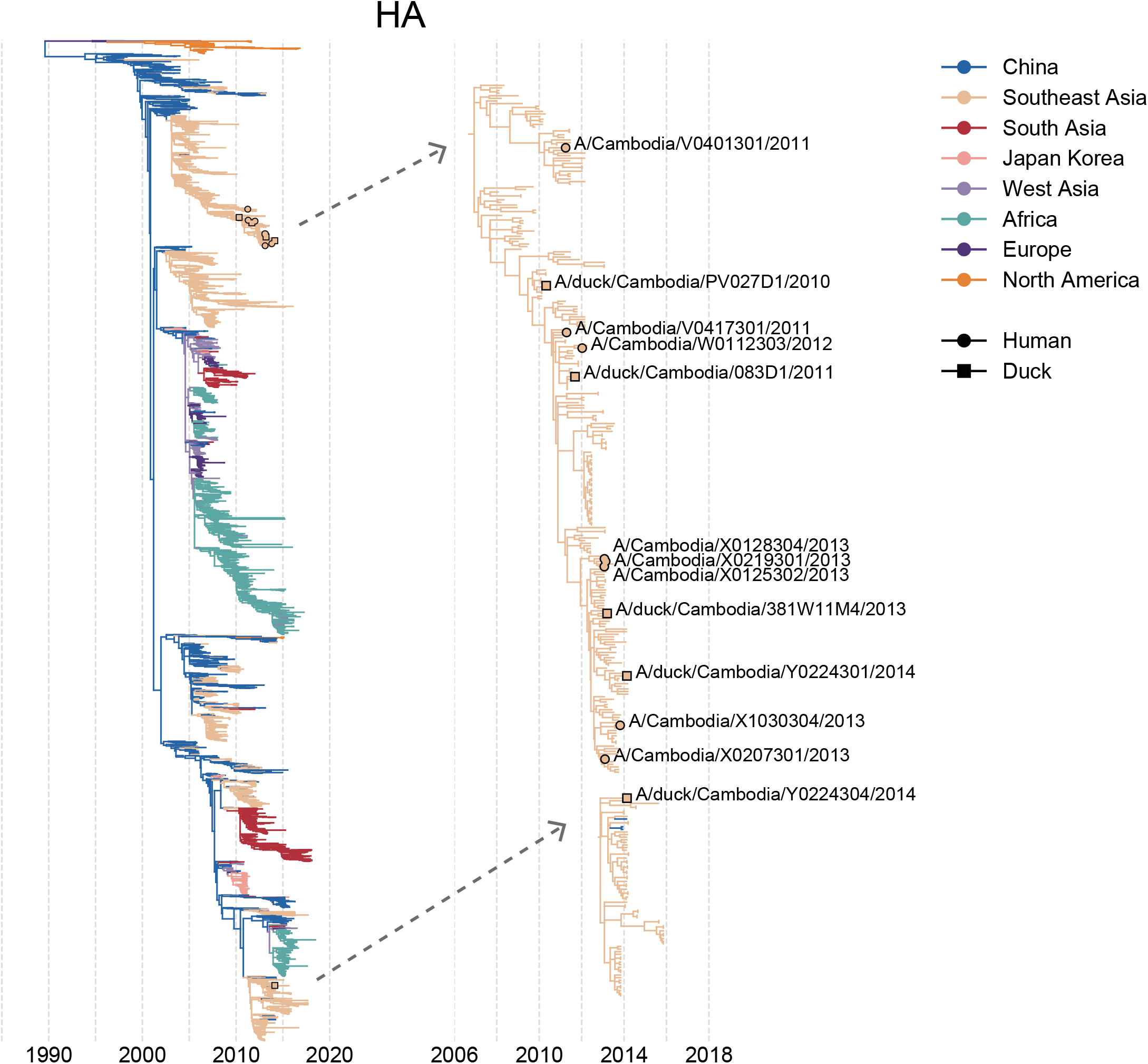
Phylogenetic placement of H5N1 samples from Cambodia. All currently available H5N1 sequences were downloaded from the Influenza Research Database and the Global Initiative on Sharing All Influenza Data and used to generate full genome phylogenies using Nextstrain’s augur pipeline as shown in the trees on the left. Phylogenies for the full genome are shown in Figure S2. Colors represent the geographic region in which the sample was collected (for tips) or the inferred geographic location (for internal nodes). The x-axis position indicates the date of sample collection (for tips) or the inferred time to the most recent common ancestor (for internal nodes). In the full phylogeny (left), H5N1 viruses from Cambodia selected for within-host analysis are indicated by tan circles with black outlines. The subtrees containing the Cambodian samples selected for within-host analysis are shown to the right and are indicated with grey, dashed arrows. In these trees, human tips are marked with a tan circle with a black outline, while duck tips are denoted with a tan square with a black outline. All samples from our within-host dataset are labelled in the subtrees with their strain name. Internal genes from samples collected prior to 2013 belong to clade 1.1.2, while internal genes from samples collected in 2013 or later belong to clade 2.3.2.1a. All HA and NA sequences in this dataset, besides A/duck/Cambodia/Y0224304/2014, belong to clade 1.1.2.

### Tajima’s *D* calculation

Tajima’s *D* was calculated with the following equation:

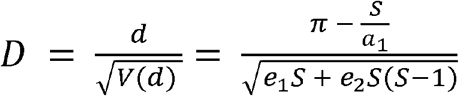

where:

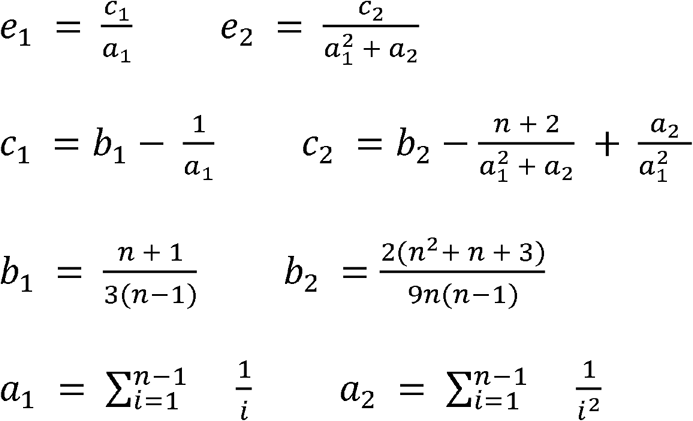

π *=* π*_N_* or π*_S_* as calculated below in “Diversity (π) calculation”, and *S* is the number of segregating sites, i.e., the number of within-host single nucleotide variants called for a given sample and coding region. Within-host variants were called as described above, requiring a minimum coverage of 100x, a minimum frequency of 1%, a minimal base quality score of Q30, and detection on both forward and reverse reads. For each sample, we treated synonymous variants and nonsynonymous variants separately, calculating *D* for nonsynonymous variation as the difference between π*_N_* and *S_N_*, and *D* for synonymous variation as the difference between π*_S_* and *S_S_*. For n, we used the average coverage across the coding region. Values shown in Fig. 2c represent mean *D* when values were combined across all human or duck samples. To calculate the 95% confidence interval, we performed a bootstrap. We resampled our *D* values with replacement, 10,000 times, and calculated the mean of the resampled values in each iteration. We then calculated the 2.5% and 97.5% percentile of these bootstrapped means and report this as the 95% confidence interval.

**Figure 2:**
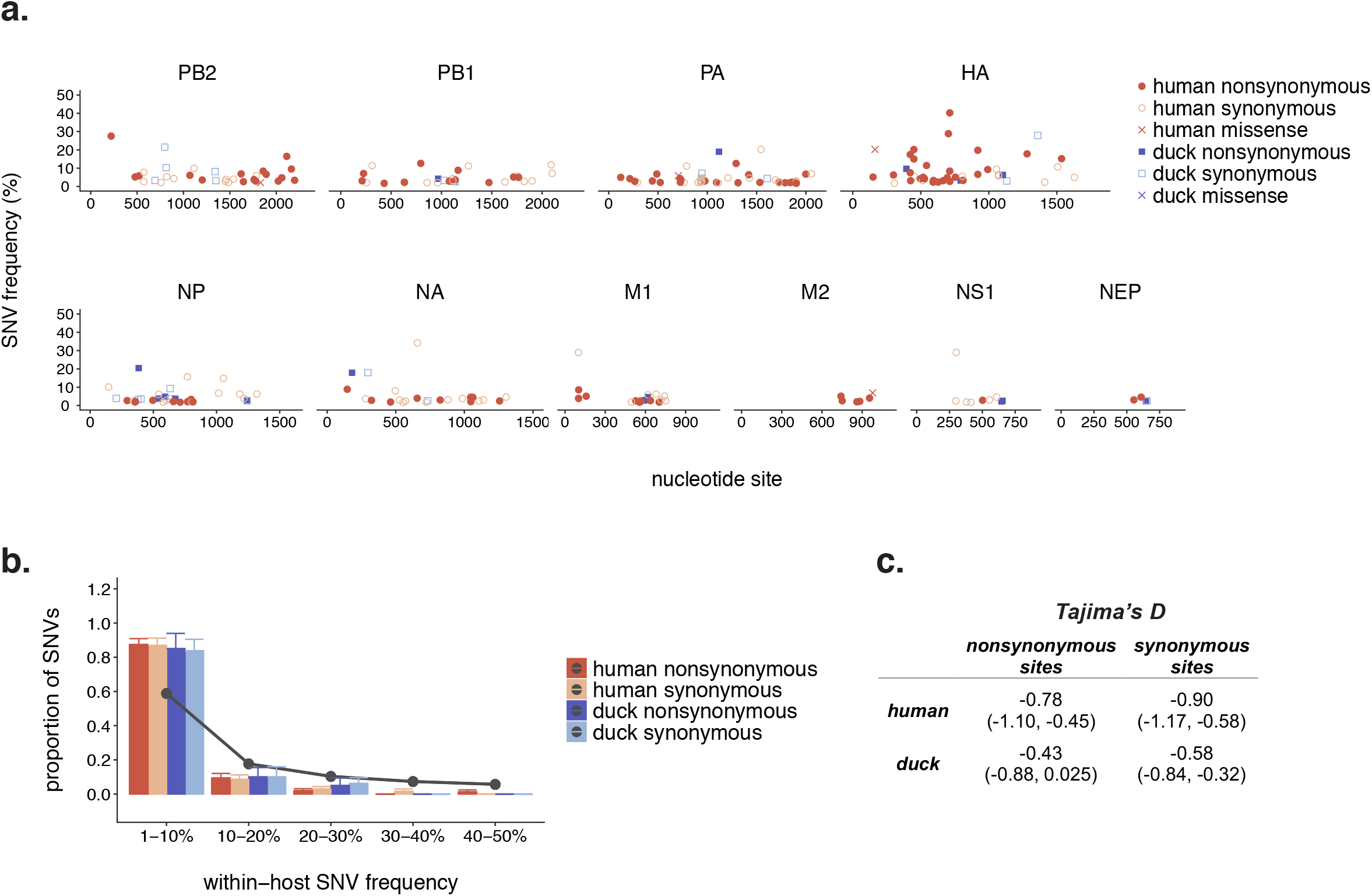
Within-host diversity in humans and ducks is dominated by low-frequency variation. **(a)** Within-host polymorphisms present in at least 1% of sequencing reads were called in all human (red) and duck (blue) samples. Each dot represents one unique single nucleotide variant (SNV), the x-axis represents the nucleotide site of the SNV, and the y-axis represents its frequency within-host. **(b)** For each sample in our dataset, we calculated the proportion of its synonymous (light blue and light red) and nonsynonymous (dark blue and dark red) within-host variants present at frequencies of 1-10%, 10-20%, 20-30%, 30-40%, and 40-50%. We then took the mean across all human (red) or duck (blue) samples. Bars represent the mean proportion of variants present in a particular frequency bin and error bars represent standard error. Grey dots and connecting lines represent the expected proportion of variants in each bin under a neutral model. **(c)** We calculated Tajima’s *D* across the full genomes of humans and ducks, separately for synonymous and nonsynonymous sites. Values represent the mean Tajima’s *D* across all humans or ducks, and values in parentheses represent the 95% confidence interval.

### Diversity (π) calculation

Within-host variants were called as described above, requiring a minimum coverage of 100x, a minimum frequency of 1%, a minimal base quality score of Q30, and detection on both forward and reverse reads. Variants were annotated as nonsynonymous or synonymous. For each sample and coding region, we computed the average number of pairwise nonsynonymous pairwise differences per nonsynonymous site (π*_N_*) and the average number of pairwise synonymous differences per synonymous site and (π*_S_*) with SNPGenie[40, 41] (https://github.com/chasewnelson/SNPGenie). We used the same set of within-host variants as reported throughout the manuscript (minimum frequency of 1%) for these diversity calculations. In both Fig. 3 and Table 2, we present the mean π*_N_* (dark colors) or π*_S_* (light colors) when values were combined across all humans (red bars) or ducks (blue bars). To calculate the standard error of these estimates, we performed a bootstrap. We resampled our diversity values with replacement, 10,000 times, and calculated the mean of the resampled values in each iteration. We then calculated the standard deviation among our sampled means, and report this as the standard error. Error bars in Fig. 3 reflect this calculated standard error.

**Figure 3:**
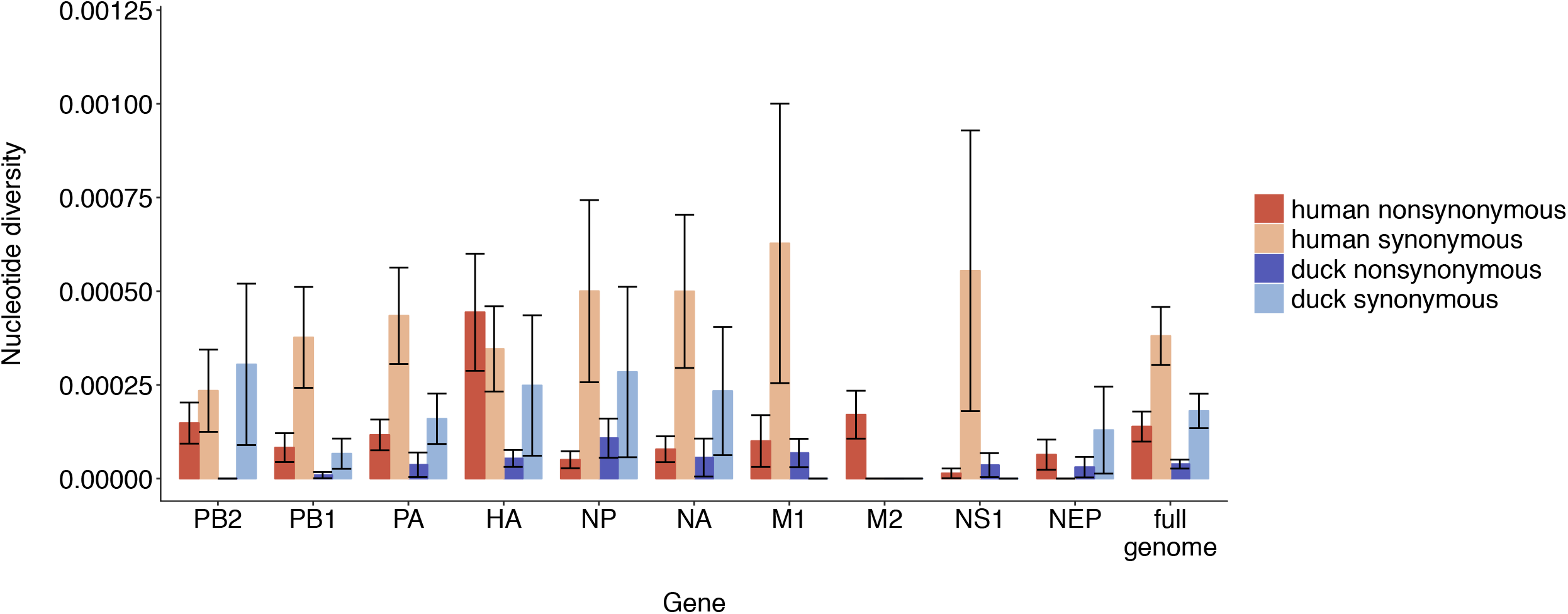
Purifying selection and genetic drift shape within-host diversity. For each sample and gene, we computed the average number of pairwise nonsynonymous differences per nonsynonymous site (π*_N_*) and the average number of pairwise synonymous differences per synonymous site (π*_S_*). We then calculated the mean for each gene and species. Each bar represents the mean and error bars represent the standard error calculated by performing 10,000 bootstrap resamplings. Human values are shown in red and duck values are shown in blue.

**Table 2:** Mean π*_N_* and π*_S_* values per gene.

| Gene | Species | Mean $\pi_N$ | Mean $\pi_S$ | $\pi_N/\pi_S$ | p-value |
| --- | --- | --- | --- | --- | --- |
| PB2 | Human | 0.00015 | 0.00023 | 0.65 | 0.50 |
| PB2 | Duck | 0.00 | 0.00031 | 0.00 | 0.27 |
| **PB1** | **Human** | **0.000083** | **0.00038** | **0.22** | **0.049** |
| PB1 | Duck | 0.000009 | 0.000066 | 0.14 | 0.31 |
| PA | Human | 0.00012 | 0.00044 | 0.27 | 0.083 |
| PA | Duck | 0.000037 | 0.00016 | 0.23 | 0.094 |
| HA | Human | 0.00044 | 0.00035 | 1.26 | 0.61 |
| HA | Duck | 0.000054 | 0.00025 | 0.22 | 0.40 |
| NP | Human | 0.000050 | 0.00050 | 0.10 | 0.12 |
| NP | Duck | 0.00011 | 0.00028 | 0.39 | 0.49 |
| NA | Human | 0.000078 | 0.0005 | 0.16 | 0.064 |
| NA | Duck | 0.000056 | 0.00023 | 0.24 | 0.27 |
| M1 | Human | 0.00010 | 0.00063 | 0.14 | 0.23 |
| M1 | Duck | 0.000068 | 0.00 | NA | 0.18 |
| **M2** | **Human** | **0.00017** | **0.00** | **NA** | **0.042** |
| M2 | Duck | 0.00 | 0.00 | NA | NA |
| NS1 | Human | 0.000014 | 0.00056 | 0.03 | 0.20 |
| NS1 | Duck | 0.000036 | 0.00 | NA | 0.37 |
| NEP | Human | 0.000064 | 0.00 | NA | 0.18 |
| NEP | Duck | 0.000030 | 0.00013 | 0.23 | 0.37 |
| **Full genome** | **Human** | **0.000139** | **0.000381** | **0.36** | **0.0059** |
| **Full genome** | **Duck** | **0.000039** | **0.00018** | **0.22** | **0.038** |

For each gene and sample, we computed nonsynonymous ( $\pi_N$ ), and synonymous ( $\pi_S$ ) diversity as the average number of pairwise differences between a set of DNA sequences.
Values of 0.00 indicate that there were no SNPs identified in that gene for that host species and mutation type. We then combined values from each sample to generate a diversity estimate for each gene and host species. Significance was assessed by a paired t-test testing the null hypothesis that $\pi_N = \pi_S$ . Bold values of $p < 0.05$ .

### Comparison to functional sites

We used the Sequence Feature Variant Types tool from the Influenza Research Database[34] to download all currently available annotations for H5 hemagglutinins, N1 neuraminidases, and all subtypes for the remaining gene segments. We then annotated each within-host SNV identified in our dataset that fell within an annotated region or site. The complete results of this annotation are available in **Table S1**. We next filtered our annotated SNVs to include only those located in sites involved in “host-specific” functions or interactions, i.e., those that are distinct between human and avian hosts. We defined host-specific functions/interactions as receptor binding, interaction with host cellular machinery, nuclear import and export, immune antagonism, 5’ cap binding, temperature sensitivity, and glycosylation. We also included sites that have been phenotypically identified as determinants of transmissibility and virulence. Sites that participate in binding interactions with other viral subunits or vRNP, conserved active site domains, drug resistance mutations, and epitope sites were not categorized as host-specific for this analysis. We annotated both synonymous and nonsynonymous mutations in our dataset, but only highlight nonsynonymous changes in Fig. 4 and Table 3.

**Figure 4:**
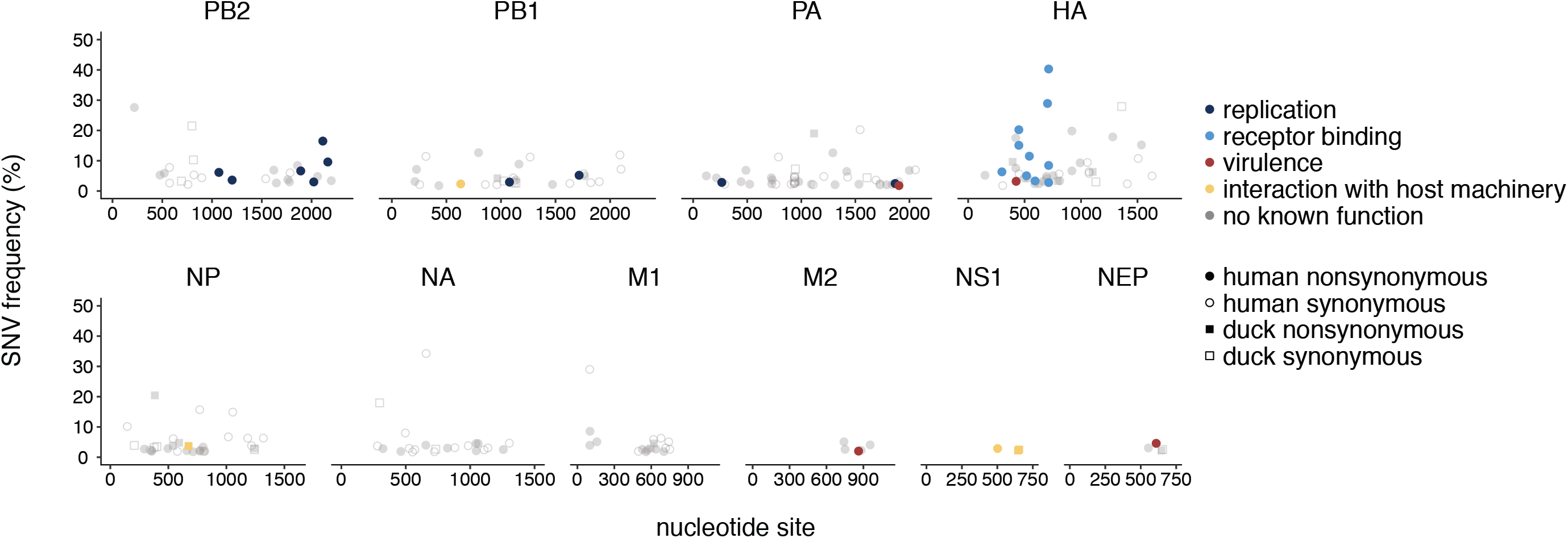
Mutations are present at functionally relevant sites. We queried each amino acid changing mutation identified in our dataset against all known annotations present in the Influenza Research Database Sequence Feature Variant Types tool. Each mutation is colored according to its function. Shape represents whether the mutation was identified in a human (circle) or duck (square) sample. Mutations shown here were detected in at least 1 human or duck sample. Filled in shapes represent nonsynonymous changes and open shapes represent synonymous mutations. Grey, transparent dots represent mutations for which no host-related function was known. Each nonsynonymous colored mutation, its frequency, and its phenotypic effect is shown in Table 3, and a full list of all mutations and their annotations are available in Table S1.

**Table 3:** Mutations identified at functionally relevant sites.

| Sample | Gene | Nt site | Ref base | Variant base | Coding region change | Freq. | Description | Type |
| --- | --- | --- | --- | --- | --- | --- | --- | --- |
| A/Cambodia/X0128304/2013 | PB2 | 1069 | A | T | N348Y | 6.15% | Putative m7GTP cap binding site[64]. | replication |
| A/Cambodia/V0401301/2011 | PB2 | 1202 | A | C | N392H | 3.61% | Putative m7GTP cap binding site[64]. | replication |
| A/Cambodia/W0112303/2012 | PB2 | 1891 | G | A | E627K | 6.63% | A Lys at 627 enhances mammalian replication[51,53]. | replication |
| A/Cambodia/X0125302/2013 | PB2 | 2022 | G | A | V667I | 2.99% | An Ile at 667 was associated with human-infecting H5N1 virus strains[65]. | replication |
| A/Cambodia/W0112303/2012 | PB2 | 2113 | A | G | N701D | 16.49% | An Asn at 701 enhances mammalian replication[55,56]. | replication |
| A/Cambodia/X0125302/2013 | PB2 | 2163 | A | G | S714G | 9.59% | An Arg at 714 enhances mammalian replication[55]. | replication |
| A/Cambodia/X1030304/2013 | PB1 | 631 | A | G | R211G | 2.34% | Nuclear localization motif. | interaction with host machinery |
| A/Cambodia/X0125302/2013 | PB1 | 1078 | A | G | K353R | 2.94% | An Arg at 353 is associated with higher replication and pathogenicity of an H1N1 pandemic strain[66]. | replication |
| A/Cambodia/X0125302/2013 | PB1 | 1716 | A | T | T566S | 5.20% | An Ala at 566 is associated with higher replication and pathogenicity of an H1N1 pandemic virus[66]. | replication |
| A/Cambodia/X0219301/2013 | PA | 265 | A | G | T85A | 2.84% | An Ile at 85 enhances polymerase activity of pandemic H1N1 in mammalian cells[67]. | replication |
| A/Cambodia/X0128304/2013 | PA | 1868 | A | G | K615R | 2.47% | An Asn at PA 615 has been associated with adaptation of avian influenza polymerases to humans[55]. | replication |
| A/Cambodia/X0207301/2013 | PA | 1903 | A | G | S631G | 1.79% | A Ser at 631 enhances virulence of H5N1 viruses in mice[68]. | virulence |
| A/Cambodia/X0128304/2013 | HA | 299 | A | G | E91G | 6.33% | A Lys at 91 enhances α-2,6 binding[43]. (H5 mature: 75) | receptor binding |
| A/Cambodia/V0417301/2011 | HA | 425 | A | G | E142G | 3.20% | Putative glycosylation site[69]. (H5 mature: 126) | virulence |
| A/Cambodia/V0401301/2011 | HA | 449 | C | T | A150V | 20.24% | A Val at 150 confers enhanced α-2,6 sialic acid binding in H5N1 viruses[58,59]. (H5 mature: 134) | receptor binding |
| A/Cambodia/X0125302/2013 | HA | 449 | C | T | A150V | 15.09% | A Val at 150 confers enhanced α-2,6 sialic acid binding in H5N1 viruses[58,59]. (H5 mature: 134) | receptor binding |
| A/Cambodia/X0128304/2013 | HA | 542 | A | C | K172T | 11.50% | Part of putative glycosylation motif that improves α-2,6 binding[70–72]. (H5 mature: 156) | receptor binding |
| A/Cambodia/V0401301/2011 | HA | 517 | T | C | Y173H | 5.04% | Residue involved in sialic acid recognition[45]. (H5 mature: 157) | receptor binding |
| A/Cambodia/V0401301/2011 | HA | 593 | A | G | N198S | 3.32% | A Lys at 198 confers α-2,6 sialic acid binding [43,73](H5 mature: 182) | receptor binding |
| A/Cambodia/X0128304/2013 | HA | 703 | A | G | T226A | 28.91% | An Ile at 226 enhanced α-2,6 sialic acid binding[63]. (H5 mature: 210) | receptor binding |
| A/Cambodia/V0401301/2011 | HA | 713 | A | T | Q238L | 2.80% | A Leu at 238 confers a switch from α-2,3 to α-2,6 sialic acid binding and is a determinant of mammalian transmission[11,12,73–76]. (H5 mature: 222) | receptor binding |
| A/Cambodia/V0417301/2011 | HA | 713 | A | T | Q238L | 8.45% | A Leu at 238 confers a switch from α-2,3 to α-2,6 sialic acid binding and is a determinant of mammalian transmission[11,12,73–76]. (H5 mature: 222) | receptor binding |
| A/Cambodia/X0125302/2013 | HA | 713 | A | G | Q238R | 40.30% | A Leu at 238 confers a switch from α-2,3 to α-2,6 sialic acid binding and is a determinant of mammalian transmission[11,12,73–76]. (H5 mature: 222) | receptor binding |
| A/duck/Cambodia/Y0224304/2014 | NP | 674 | C | T | T215I | 3.69% | Nuclear targeting motif[77]. | interaction with host machinery |
| A/Cambodia/X1030304/2013 | M2 | 861 | G | A | C50Y | 2.03% | A Cys at position 50 is a palmitoylation site that enhances virulence[78,79]. | virulence |
| A/Cambodia/X0128304/2013 | NS1 | 502 | C | T | P159L | 2.8% | Part of the NS1 nuclear export signal mask[80]. | interaction with host machinery |
| A/duck/Cambodia/Y0224301/2014 | NS1 | 646 | T | C | L207P | 2.22% | NS1 flexible tail, which interacts with host machinery[81]. | interaction with host machinery |
| A/duck/Cambodia/Y0224301/2014 | NS1 | 654 | C | T | P210S | 2.55% | NS1 flexible tail, which interacts with host machinery[81]. | interaction with host machinery |
| A/Cambodia/X0207301/2013 | NEP | 609 | A | G | E47G | 4.59% | This site was implicated in enhanced virulence of H5N1 viruses in ferrets[82]. | virulence |

### Shared sites permutation test

To test whether human or duck samples shared more polymorphisms than expected by chance, we performed a permutation test. We first counted the number of variable amino acid sites, *n*, in which an SNV altered the coded amino acid, across coding regions and samples. For example, if two SNVs occurred in the same codon site, we counted this as 1 variable amino acid site. Next, for each gene and sample, we calculated the number of amino acid sites that were covered with sufficient sequencing depth that a mutation could have been called using our SNV calling criteria. To do this, we calculated the length in amino acids of each coding region, *L*, that was covered by at least 100 reads. Non-coding regions were not included. For each coding region and sample, we then simulated the effect of having *n* variable amino acid sites placed randomly along the coding region between sites 1 to *L*, and recorded the site where the polymorphism was placed. After simulating this for each gene and sample, we counted the number of sites that were shared between at least 2 human or at least 2 duck samples. This process was repeated 100,000 times. The number of shared polymorphisms at each iteration was used to generate a null distribution, as shown in Fig. 5b. We calculated p-values as the number of iterations for which there were at least as many shared sites as observed in our actual data, divided by 100,000. For the simulations displayed in Fig. 5c and Fig. 5d, we wanted to simulate the effect of genomic constraint, meaning that only some fraction of the genome could tolerate mutation. For these analyses, simulations were done exactly the same, except that the number of sites at which a mutation could occur was reduced to 70% (Fig. 5c) or 60% (Fig. 5d). Code for performing the shared sites permutation test is freely available at https://github.com/blab/h5n1-cambodia/blob/master/figures/figure-5b-shared-sites-permutation-test.ipynb.

**Figure 5:**
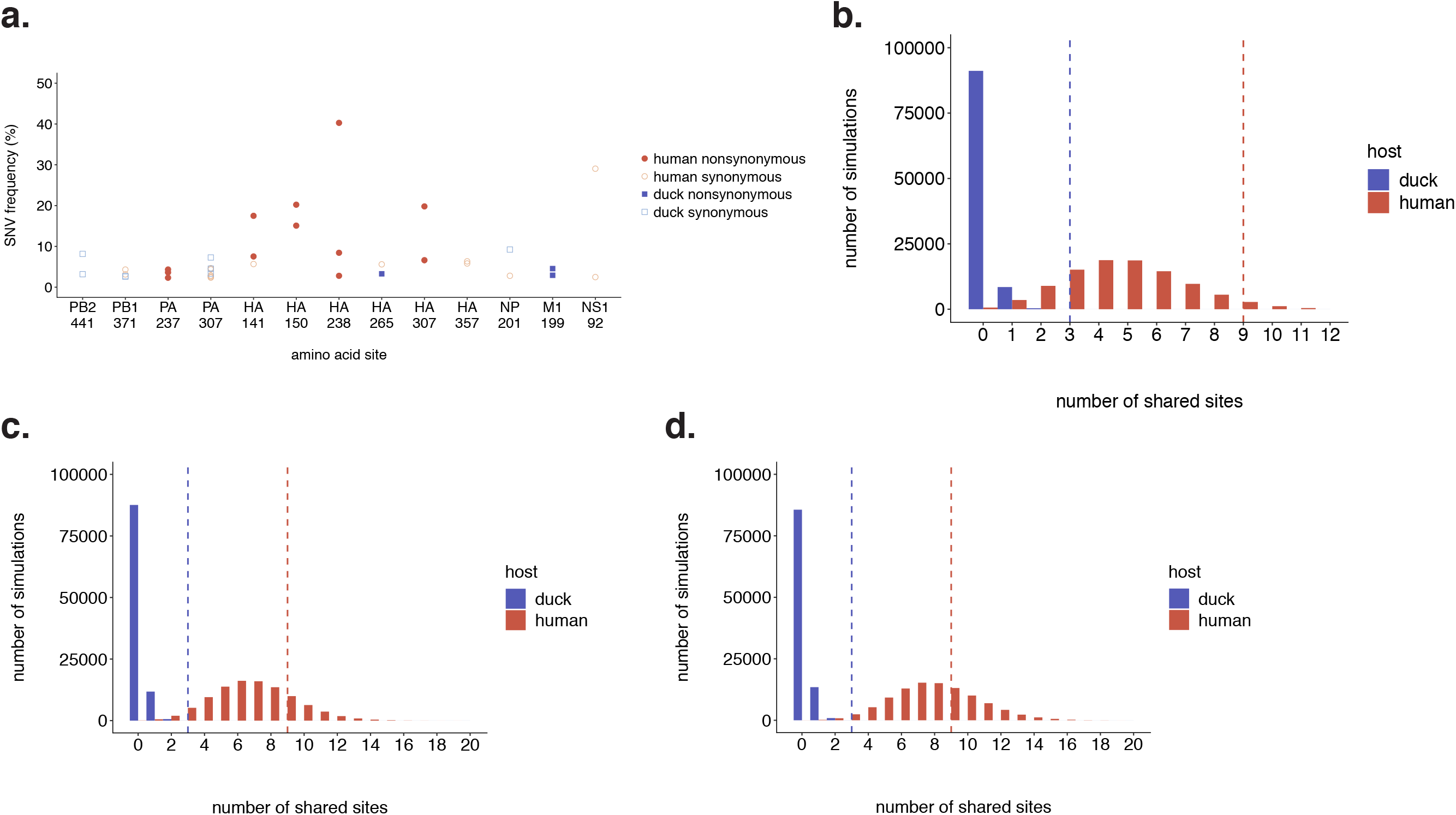
Ducks share more polymorphisms than expected by chance. **(a)** All amino acid sites that were polymorphic in at least 2 samples are shown. This includes sites at which each sample had a polymorphism at the same site, but encoded different variant amino acids. There are 3 amino acid sites that are shared by at least 2 duck samples, and 9 polymorphic sites shared by at least 2 human samples. 3 synonymous changes are detected in both human and duck samples (PB1 371, PA 397, and NP 201). Frequency is shown on the y-axis. **(b)** To test whether the level of sharing we observed was more or less than expected by chance, we performed a permutation test. The x-axis represents the number of sites shared by at least 2 ducks (blue) or at least 2 humans (red), and the bar height represents the number of simulations in which that number of shared sites occurred. Actual observed number of shared sites (3 and 9) are shown with a dashed line. **(c)** The same permutation test as shown in **(b)**, except that only 70% of amino acid sites were permitted to mutate. **(d)** The same permutation test as shown in **(b)**, except that only 60% of amino acid sites were permitted to mutate.

### Reconstruction of host transitions along the phylogeny

We used the phylogenetic trees in **Fig. S2** to infer host transitions along each gene’s phylogeny. As described above, we used TreeTime[39] to reconstruct ancestral nucleotide states at each internal node and infer amino acid mutations along each branch along these phylogenetic trees. We then classified host transition mutations along branches that lead to human or avian tips as follows (Fig. 6a). For each branch in the phylogeny, we enumerated all tips descending from that branch. If all descendent tips were human, we considered this a monophyletic human clade. If the current branch’s ancestral node also led to only human descendants, we labelled the current branch a “to-human” branch. If a branch leading to a monophyletic human clade had an ancestral node that included avian and human descendants, then we considered the current branch an “avian-to-human” branch, and also labelled it as “to-human”. All other branches were considered “to-avian” branches. We did not explicitly allow for human-to-avian branches in this analysis. Because avian sampling is poor relative to human sampling, and because H5N1 virus circulation is thought to be maintained by transmission in birds, we chose to only label branches explicitly leading to human infections as to-human branches. We also reasoned that for instances in which a human tip appears to be ancestral to an avian clade, this more likely results from poor avian sampling than from true human-to-avian transmission. Using these criteria, we then gathered the inferred amino acid mutations that occurred along each branch in the phylogeny, and counted the number of times they were associated with each type of host transition. We then queried each SNV detected within-host in our dataset, in both human and duck samples, to determine the number of host transitions that they occurred on in the phylogeny, as shown in Fig. 6b. To test whether individual mutations were enriched along branches leading to human infections, we performed Fisher’s exact tests comparing the number of to-avian and to-human transitions along which the mutation was detected vs. the overall number of to-avian and to-human transitions that were observed along the tree. Mutations that showed statistically significant enrichment are annotated in Fig. 6b.

**Figure 6:**
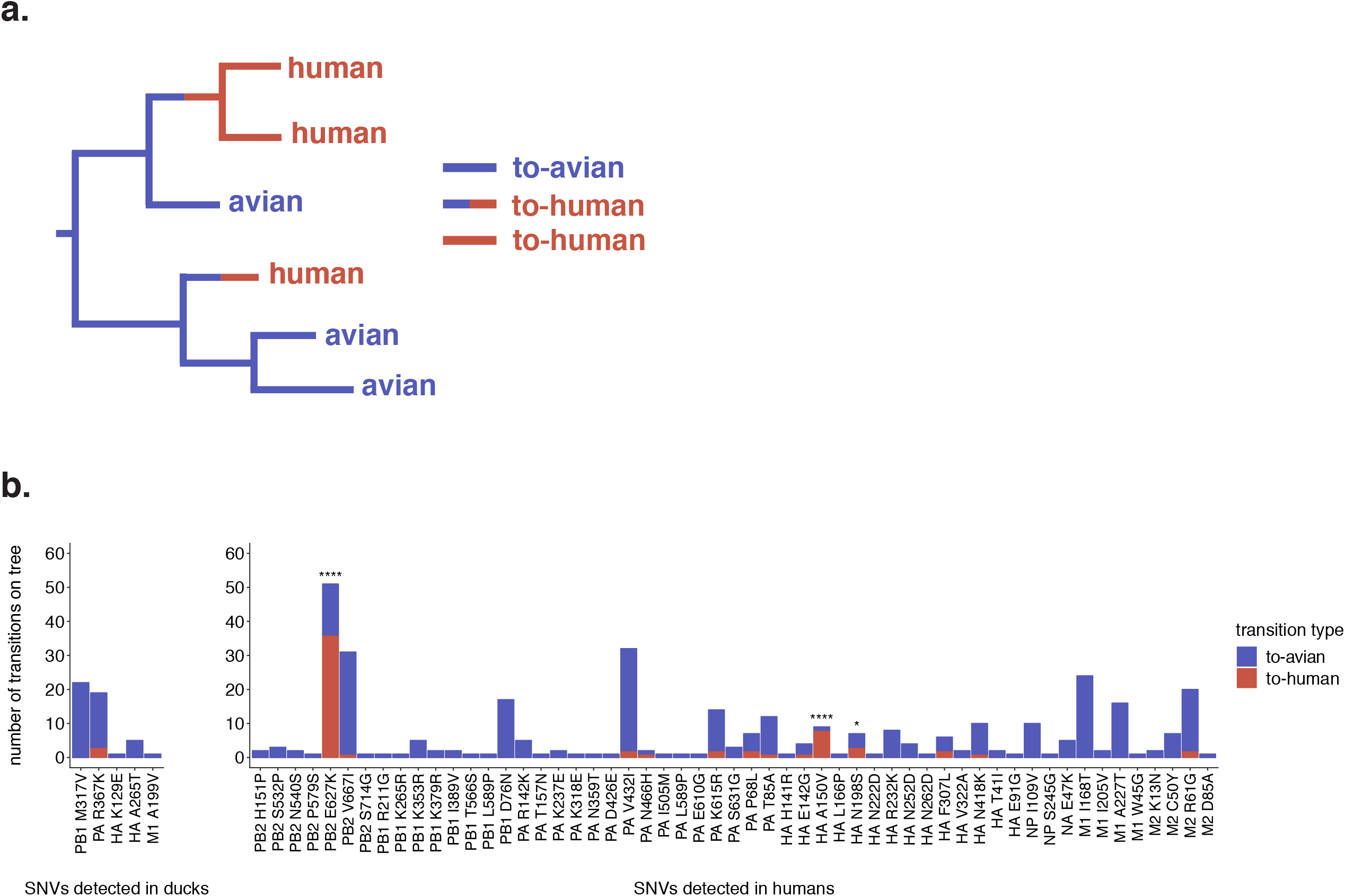
A small subset of within-host variants are enriched on spillover branches. **(a)** A schematic for how we classified host transitions along the phylogeny. Branches within monophyletic human clades were labelled “to-human” (red branches). Branches leading to a monophyletic human clade, whose parent node had avian children were also labelled as “to-human” (half red, half blue branches), and all other branches were labelled “to-avian” (blue branches). **(b)** Each amino acid-changing SNV we detected within-host in either ducks (left) or humans (right) that was present in the H5N1 phylogeny is displayed. Each bar represents an amino acid mutation, and its height represents the number of to-avian (blue) or to-human (red) transitions in which this mutation was present along the H5N1 phylogeny. Significance was assessed with a Fisher’s exact test. * indicates p < 0.05, **** indicates p < 0.0001.

### General availability of analysis software and data

All code used to analyze data and generate figures for this manuscript are publicly available at https://github.com/blab/h5n1-cambodia. Raw FASTQ files with human reads removed are available under SRA accession number PRJNA547644, and accessions SRX5984186-SRX5984198. All reported variant calls and phylogenetic trees are available at https://github.com/blab/h5n1-cambodia/tree/master/data.

## Results

### Sample selection and dataset information

We analyzed full-genome sequence data from primary, influenza virus-confirmed samples from infected humans and domestic ducks from Cambodia (Table 1). Four domestic duck samples (pooled organs) were collected as part of poultry outbreak investigations, while one was collected during live bird market surveillance (pooled throat and cloacal swab)[4]. All human samples (throat swabs) were collected via event-based surveillance upon admittance to various hospitals throughout Cambodia[22]. Because of limited sample availability and long storage times, generating duplicate sequence data for each sample was not possible. We therefore focused on samples whose viral RNA copy numbers after viral RNA extraction were ≥10^3^ copies/μl of buffer as assessed by RT-qPCR (Table 1), and whose mean coverage depth exceeded 100x (**Fig. S1**). We analyzed full genome data for 7 human and 5 duck samples, and near complete genome data for A/Cambodia/X0128304/2013, for which we lack data from the PB1 gene.

H5 viruses circulating in Cambodia were exclusively clade 1.1.2[4] until 2013, when a novel reassortant virus emerged[42]. This reassortant virus expressed a hemagglutinin (HA) and neuraminidase (NA) from clade 1.1.2, with internal genes from clade 2.3.2.1a[22]. All 2013/2014 samples in our dataset come from this outbreak, while samples collected prior to 2013 are clade 1.1.2 (Table 1, Fig. 1, and **Fig. S2**). All HA sequences (with the exception of A/duck/Cambodia/Y0224304/2014, which expresses a divergent HA) derive from the same lineage that has been circulating in southeast Asia for years (Fig. 1). For the internal gene segments, samples collected between 2010-2012 and samples collected between 2013-2014 fall into distinct parts of the tree, each nested within the diversity of other southeast Asian viruses (**Fig. S2**). The 2013 reassortant viruses share 4 amino acid substitutions in HA, S123P, S133A, S155N, and K266R[22] (H5, mature peptide numbering). S133A and S155N have been linked to improved α-2,6 linked sialic acid binding, independently and in combination with S123P[43–45]. All samples encode a polybasic cleavage site in HA (XRRKRR) between amino acids 325-330 (H5, mature peptide numbering), a virulence determinant for H5N1 AIVs[46, 47], and a 20 amino acid deletion in NA. This NA deletion is a well-documented host range determinant[48–51].

Duck samples are not immediately ancestral to the human samples in our dataset, and they therefore are unlikely to represent transmission pairs. We therefore treat these samples as examples of within-host diversity in naturally infected humans and ducks, rather than direct transmission pairs. With this caveat, we aimed to use this subset of 8 human and 5 duck samples to determine whether positive selection would promote adaptation in humans. Positive selection increases the frequency of beneficial variants, and is often identified by tracking mutations’ frequencies over time. While multiple time points were not available in our dataset, all human samples were collected 5-12 days after reported symptom onset[22]. Animal infection studies have observed drastic changes in within-host variant frequencies in 3-7 days[11, 13], suggesting that 5-12 days post symptom onset may provide sufficient time for transmitted diversity to be altered within-host. We reasoned that while we expect positive selection to promote the emergence of human-adapting mutation in humans, H5N1 viruses should be well-adapted for replication in ducks, which are a natural host species. We therefore hypothesize to observe the following patterns: (1) During replication in humans, positive selection should increase the frequencies of human-adaptive mutations, resulting in elevated rates of nonsynonymous diversity and a higher proportion of high-frequency variants. In contrast, viruses in ducks should be fit for replication and be shaped by purifying selection, leading to an excess of synonymous variation and an excess of low-frequency variants. (2) Viruses in humans should harbor mutations phenotypically linked to mammalian adaptation. (3) If selection is strong at a particular site, then viruses in humans should exhibit evidence for convergent evolution, i.e., the same mutation arising across multiple samples. (4) If human-adaptive variants arising within humans are present on the H5N1 phylogeny, then they should be more likely to occur on branches leading to human infections than on branches leading to bird infections.

### Within-host diversity in humans and ducks is dominated by low-frequency variation

We called within-host variants across the genome that were present in ≥1% of sequencing reads and occurred at a site with a minimum read depth of 100x and a minimum PHRED quality score of 30 (see Methods for details). All coding region changes are reported using native H5 numbering, including the signal peptide, unless otherwise noted. Most single nucleotide variants (SNVs) were present at low frequencies (Fig. 2a). We identified a total of 206 SNVs in humans (111 nonsynonymous, 91 synonymous, 4 missense) and 40 in ducks (16 nonsynonymous, 23 synonymous, 1 missense). Human samples had more SNVs than duck samples on average (mean SNVs per sample: humans = 26 ± 19, ducks = 8 ± 3, p = 2.79 x 10^-17^, Fisher’s exact test), although the number of SNVs per sample was variable among samples in both species (**Fig. S3**).

To determine whether humans had more high-frequency variants than ducks, we generated a site frequency spectrum (Fig. 2b). Purifying selection removes new variants from the population, generating an excess of low-frequency variants, while positive selection promotes accumulation of high-frequency polymorphisms. Exponential population expansion also leads to an excess of low-frequency variation. In both humans and ducks, over 80% of variants (both synonymous and nonsynonymous) were present in <10% of the population, and the distribution of SNV frequencies were strikingly similar (Fig. 2b). In both host species, there is an excess of low-frequency variation compared to the expectation under a neutral model (no population size changes or selection), and a deficiency of intermediate and high-frequency variants (Fig. 2b, grey dots and connecting line). Overall, the frequencies of SNVs in humans and ducks were not statistically different (p=0.11, Mann Whitney U test), and mean SNV frequencies were similar (mean SNV frequency in human samples = 5.8%, mean in duck samples = 6.6%).

To determine whether the excess of low-frequency variation we observed was better explained by purifying selection or demography, we summarized the frequency spectrum by calculating Tajima’s *D* (Fig. 2c). Tajima’s *D* measures the difference between the average number of pairwise differences between a set of sequences (π) with the number of variable sites (*S*). π is weighted by variant frequencies, and will be largest when the population has a large number of high-frequency variants, while *S* is simply a count of the number of variable sites, and is not impacted by variant frequencies. Both population expansion and purifying selection should lead to an excess of low-frequency variation and negative Tajima’s *D*. However, while population expansion should impact nonsynonymous and synonymous sites equally, purifying selection should have a greater effect on nonsynonymous variants. If the excess of low-frequency variation we observed was driven solely by demographic factors, then we expect synonymous and nonsynonymous sites to have similar Tajima’s *D* values, while purifying selection should lead to more negative Tajima’s *D* values at nonsynonymous sites. When calculated across the full genome, Tajima’s *D* was similar between humans and ducks, and was comparable when calculated for synonymous and nonsynonymous sites. Taken together, these data suggest that in both humans and ducks, viral populations are dominated by low-frequency variation. Furthermore, this excess of low-frequency variation can be explained by within-host population expansion.

### Purifying selection and genetic drift shape within-host diversity

Comparing nonsynonymous (π*_N_*) and synonymous (π*_S_*) polymorphism in a population is another common measure for selection that is robust to differences in sequencing coverage depth[52]. An excess of synonymous polymorphism (π*_N_*/π*_S_* < 1) indicates purifying selection, an excess of nonsynonymous variation (π*_N_*/π*_S_* > 1) suggests positive selection, and approximately equal rates (π*_N_*/π*_S_* ∼ 1) suggest that genetic drift is the predominant force shaping diversity. We calculated the average number of nonsynonymous and synonymous pairwise differences between DNA sequences, and normalized these values to the number of synonymous and nonsynonymous sites. In both species, most genes exhibited π*_N_* < π*_S_*, although there was substantial variation among samples (Table 2 and Fig. 3). The difference between π*_s_* and π*_N_* was generally not statistically significant (Table 2), with the exception of human M2 (π*_N_* = 0.00017, π*_S_* = 0, p = 0.042, paired t-test) and PB1 (π*_N_* = 0.000083, π*_S_* = 0.00038, p = 0.049, paired t-test), which exhibited weak evidence of purifying selection. When diversity estimates across all genes were combined, both species exhibited π*_N_*/π*_S_* < 1 (Fig. 3) (human π*_N_*/π*_S_*= 0.36, p = 0.0059, unpaired t-test; duck π*_N_*/π*_S_*= 0.21, p = 0.038, unpaired t-test). Genome-wide diversity was not correlated with days post symptom onset (**Fig. S4a**). Taken together, these data suggest that H5N1 within-host populations in both humans and ducks are broadly shaped by weak purifying selection and genetic drift. We do not find evidence for widespread positive selection in any individual coding region.

### SNVs are identified in humans at functionally relevant sites

Influenza phenotypes can be drastically altered by single amino acid changes. We took advantage of the Influenza Research Database^29^ Sequence Feature Variant Types tool, a catalogue of amino acids that are critical to protein structure and function, and that have been experimentally linked to functional alteration. We downloaded all available annotations for H5 HAs, N1 NAs, and all subtypes for the remaining proteins, and annotated each mutation in our dataset that fell within an annotated region (**Table S1**). We then filtered these annotated amino acids to include only those located in sites involved in host-specific functions (see Methods for details).

Of the 218 unique, polymorphic amino acid sites in our dataset (including both human and duck samples), we identified 34 nonsynonymous mutations at sites involved in viral replication, receptor binding, virulence, and interaction with host cell machinery (Fig. 4). Some sites are explicitly linked to H5N1 virus mammalian adaptation (Table 3). PB2 E627K was detected as a minor variant in A/Cambodia/W0112303/2012, and in A/Cambodia/V0417301/2011 at consensus. A lysine at position 627 is a conserved marker of human adaptation[51, 53] that enhances H5N1 replication in mammals[11,12,51,54]. A/Cambodia/W0112303/2012 also encoded PB2 D701N at consensus. Curiously, this patient also harbored the reversion mutation, N701D, at low-frequency within-host. An asparagine (N) at PB2 701 enhances viral replication and transmission in mammals[55, 56], while an aspartate (D) is commonly identified in birds. We cannot distinguish whether the founding virus harbored an asparagine or aspartate, so our data are consistent with two possibilities: transmission of a virus harboring asparagine and within-host generation of aspartate; or, transmission of a virus with asparate followed by within-host selection but incomplete fixation of asparagine. All other human and avian samples in our dataset encoded the “avian-like” amino acids, glutamate at PB2 627, and aspartate at PB2 701. None of the adaptive polymerase mutations that were recently identified by Welkers et al.[17] in H5N1 virus-infected humans in Indonesia were present in our samples, nor were any of the human-adaptive mutations identified in a recent deep mutational scan of PB2[57].

We also identified HA mutations linked to human receptor binding. Two human samples encoded an HA A150V mutation (134 in mature, H5 peptide numbering, Fig. 4). A valine at HA 150 improves α-2,6 linked sialic acid binding in H5N1 viruses[58, 59], and was also identified in H5N1 virus-infected humans in Vietnam[16]. Finally, HA Q238L was detected in A/Cambodia/V0417301/2011 and A/Cambodia/V0401301/2011. HA 238L (222 in mature, H5 peptide numbering) was shown in H5N1 virus transmission studies to confer a switch from α-2,3 to α-2,6 linked sialic acid binding[11] and mediate transmission[11, 12]. An HA Q238R mutation was identified in A/Cambodia/X0125302/2013, although nothing is known regarding an arganine (R) at this site.

Mutations annotated as host-specific were not detected at higher frequencies than non-host-specific mutations (mean frequency for host-specific mutations = 8.2% ± 8.8%, mean frequency for non-host-specific mutations = 5.2% ± 4.7%, p-value = 0.084, unpaired t-test). Additionally, the proportion of mutations that were host-specific was not higher in samples from longer infections (p-value = 0.72, Fisher’s exact test, **Fig. S4b**). All 8 human samples harbored at least 1 mutant in a host-specific site. Critically though, the functional impacts of influenza virus mutations strongly depend on sequence context[60], and we did not phenotypically test these mutations. We caution that confirming functional impacts for these mutations would require further study. Still, our data show that putative human-adapting mutations are generated during natural spillover. Our results also highlight that even mutations that have been predicted to be strongly beneficial (e.g., PB2 627K and HA 238L) may remain at low frequencies in vivo.

### Shared diversity is limited

Each human H5N1 infection is thought to represent a unique avian spillover event. If selection is strong at a given site in the genome, then mutations may arise at that site independently across multiple patients. We identified 13 amino acid sites in our dataset that were polymorphic in at least 2 samples, 4 of which were detected in both species (PB1 371, PA 307, HA 265 and NP 201). Of the 34 unique polymorphic amino acid sites in ducks, 3 sites were shared by at least 2 duck samples; of the 188 unique polymorphic amino acid sites in humans, 9 were shared by at least 2 human samples (Fig. 5a). Two of these shared sites, HA 150 and HA 238, are linked to human-adapting phenotypes (Table 3). To determine whether the number of shared sites we observe is more or less than expected by chance, we performed a permutation test. For each species, we simulated datasets with the same number of sequences and amino acid polymorphisms as our actual dataset, but assigned each polymorphism to a random amino acid site. For each iteration, we then counted the number of polymorphic sites that were shared by ≥2 samples. We ran this simulation for 100,000 iterations for each species, and used the number of shared sites per iteration to generate a null distribution (Fig. 5b, colored bars). Comparison to the observed number of shared sites (3 and 9, dashed lines in Fig. 5b), confirmed that humans share slightly more polymorphisms than expected by chance (p = 0.046), while ducks share significantly more (p = 0.00006).

Viral genomes are highly constrained [61], which could account for the convergence we observe. Experimental measurements of the distribution of fitness effects in influenza A virus have estimated that approximately 30% of genome mutations are lethal [61], while estimates from other RNA viruses suggest that lethal percentage ranges from 20-40% [62]. We repeated our simulations to restrict the number of amino acid sites that could tolerate a mutation to 70% or 60%, representing a lethal fraction of 30% or 40%. When 70% of the coding region was permitted to mutate, ∼23% of simulations resulted in ≥9 shared sites in humans (p = 0.23)(Fig. 5c), and when 60% of the genome was permitted to mutate, ∼39% of simulations resulted in ≥9 shared sites (p = 0.39)(Fig. 5d). In contrast, the probability of observing 3 shared sites among duck samples remained low regardless of genome constraint (70% of genome tolerates mutation: p = 0.00014; 60% of genome tolerates mutation: p = 0.00028), suggesting a significant, although low, level of convergence (Fig. 5c, d). Taken together, our results suggest that duck samples share significantly more variants than expected by chance. In humans, despite the presence of shared polymorphisms with known human-adaptive phenotypes, the degree of convergence we observe is no more than expected given genome constraint.

### Within-host SNVs are not enriched on spillover branches

If within-host mutations are human-adapting, then those mutations should be enriched among H5N1 viruses that have caused human infections in the past. To test this hypothesis, we inferred full genome phylogenies using all available full-genome H5N1 viruses from the EpiFlu[32, 33] and IRD[34] databases (Fig. 1 and **Fig. S2)**, reconstructed ancestral nucleotide states at each internal node, and inferred amino acid mutations along each branch. We then classified host transition mutations along branches that led to human or avian tips (Fig. 6a). If a branch fell within a clade that included only human tips, that branch was labelled as a “to-human” transition. If a branch led to a human-only clade but its ancestral branch included avian descendants, this was inferred to be an avian-to-human transition, and was also labelled as “to-human”. All other transitions were labelled “to-avian” (Fig. 6a, see Methods for more details). We then curated the mutations that occurred on each type of host transition, and compared these counts to the mutations identified within-host in our dataset.

Of the 120 nonsynonymous within-host SNVs we identified in our dataset, 60 (50%) were not detected on the phylogeny at all. This suggests that many of the mutations generated within-host are purged from the H5N1 viral population over time. Additionally, because humans are generally dead-end hosts for H5N1 viruses, even human-adapting variants arising within-host are likely to be lost due to lack of onward transmission. Of the within-host mutations that were detected on the phylogeny, most occurred on branches leading to avian infections (Fig 6b, blue bars). However, there were a few exceptions (Fig 6b, red bars). Across the phylogeny, we enumerated a total of 31,939 to-avian transitions, and 2,787 to-human transitions, so that we expect a 11.46:1 ratio of to-avian transitions relative to to-human transitions. In contrast, PB2 E627K was heavily enriched among human infections, detected on 15 to-avian transitions and 36 to-human transitions (p = 4.21 x 10^-28^, Fisher’s exact test). HA A150V was detected in only one to-avian transition, but in 8 to-human transitions (p = 1.46 x 10^-8^, Fisher’s exact test), and HA N198S was detected on 4 to-avian transitions and 3 to-human transitions (p = 0.014, Fisher’s exact test). Although nothing is known regarding a serine at HA 198, a lysine at that site can confer α-2,6-linked sialic acid binding[43, 63]. Taken together, these data suggest that the majority of mutations detected within-host are not associated with human spillover. However, they agree with selection for human-adapting phenotypes at a small subset of sites (PB2 E627K, HA A150V, HA N198S).

## Discussion

Our study utilizes a unique dataset of to quantify H5N1 virus diversity in natural spillover infections. We establish a set of hypotheses to interrogate whether H5N1 viruses adapt to humans during natural spillover, and find support for two of them. We detect putative human-adapting mutations (PB2 E627K, HA A150V, and HA Q238L) during human infection, two of which arose multiple times (supporting hypothesis 2). PB2 E627K and HA A150V are enriched along phylogenetic branches leading to human infections, supporting their potential role in human adaptation (supporting hypothesis 4). However, we also find that population growth, genetic drift, and weak purifying selection broadly shape viral diversity in both hosts (rejecting hypothesis 1), and that convergent evolution in human viruses can be explained by genomic constraint (rejecting hypothesis 3). Together, our data show that during spillover, H5N1 viruses have the capacity to generate well-known markers of mammalian adaptation in multiple, independent hosts. However, none of these markers reached high-frequencies within-host. We speculate that during spillover, short infection times, genetic drift, demography, and purifying selection may together limit the capacity of H5N1 viruses to evolve extensively during a single human infection.

Although data from spillovers are limited, our results align with data from Vietnam[16] and Indonesia[17]. Welkers et al.[17] identified markers of mammalian replication (PB2 627K) and transmission (HA 220K) in humans, but found that adaptive markers were not widespread. Welkers et al. also characterized new mutations that improved human replication, suggesting that there are yet undiscovered pathways for adaptation. Imai et al.[16] characterized SNVs in H5N1-infected humans that altered viral replication, receptor binding, and interferon antagonism, but these mutations stayed at low frequencies. Imai et al. also showed that most within-host variants elicited neutral or deleterious effects on protein function in humans, aligning with the purifying selection we detect within-host, and the absence of ∼50% of within-host variants in the phylogeny. These findings also agree with predictions by Russell et al.[14], who hypothesized that H5N1 viruses would generate human-adapting mutations during infection, but that these mutations would remain at low frequencies and fail to be transmitted.

One unexpected result is that mutations hypothesized to be strongly beneficial, like PB2 627K and HA 238L, remained low-frequency during infection. These mutations could have arisen late in infection or been linked to deleterious mutations. Additionally, epistasis is crucial to influenza virus evolution, and mutations that promote human adaptation in one background may not be well-tolerated in others. PB2 E627K is widespread among clade 2.2.1 H5N1 viruses, but only sparsely detected in other H5N1 clades. Soh et al.[57] recently uncovered strongly human-adapting PB2 mutations that are rare in nature, likely because they are inaccessible via single site mutations. Genetic background plays a vital role in determining how AIVs evolve, and may at least partially explain our findings. Importantly, our study involves a small number of samples from a single geographic location, and two H5N1 virus clades. Continued characterization of H5N1 virus spillover in other clades is necessary to define whether our observations are generalizable across H5N1 virus outbreaks.

An important caveat of our study is that the human and duck samples described likely do not represent transmission pairs. Although the samples analyzed in this study descend from the same HA lineage (with the exception of A/duck/Cambodia/Y0224304/2014), the duck samples are not phylogenetically ancestral to the human samples in this dataset (Fig. 1 and **Fig. S2**), and most likely were not the source of the human infections. We therefore caution that each sample in this dataset merely represents an example of within-host diversity in a naturally infected host, rather than a before and after snapshot of individual cross-species transmission events.

Assessing zoonotic risk is critical but challenging. By quantifying patterns of within-host diversity, identifying mutations at adaptive sites, measuring convergent evolution, and comparing within-host diversity to long-term evolution, we can assemble a nuanced understanding of AIV evolution. These methods provide a foundation for understanding cross-species transmission that can readily be applied to other avian influenza virus datasets, as well as newly emerging zoonotic viruses.

## Supporting information

combined supplemental figures

## Acknowledgments

We would like to thank Katherine Xue for her careful reading and comments on the manuscript.

## Supporting information legends

**Figure S1: Genome coverage**

The mean coverage depth at each nucleotide site (x-axis) for each gene across our 8 human and 5 duck samples is shown. Solid black lines represent the mean coverage across samples, and the grey shaded area represents the standard deviation of coverage depth across samples.

**Figure S2: Full genome phylogenetic placement of H5N1 virus samples from Cambodia**

All currently available H5N1 virus sequences were downloaded from the Influenza Research Database and the Global Initiative on Sharing All Influenza Data and used to generate full genome phylogenies using Nextstrain’s augur pipeline. Colors represent the geographic region in which the sample was collected (for tips) or the inferred geographic location (for internal nodes). The x-axis position indicates the date of sample collection (for tips) or the inferred time to the most recent common ancestor (for internal nodes). In the full phylogenies (left), H5N1 viruses from Cambodia selected for within-host analysis are indicated by tan circles with black outlines. The subtrees containing the Cambodian samples selected for within-host analysis are shown to the right in the order that they appear in the full tree. In these trees, human tips are marked with a tan circle with a black outline, while duck tips are denoted with a tan square with a black outline. Both human and duck tips are labelled with their strain names. Internal genes from samples collected prior to 2013 belong to clade 1.1.2, while internal genes from samples collected in 2013 or later belong to clade 2.3.2.1a. All HA and NA sequences in this dataset, besides A/duck/Cambodia/Y0224304/2014, belong to clade 1.1.2.

**Figure S3: All within-host variants detected in our dataset**

All within-host variants detected in our study are shown. Each row represents one sample and each column represents one gene. The x-axis shows the nucleotide site and the y-axis shows the frequency that the variant was detected within-host. Filled circles represent nonsynonymous changes, while open circles represent synonymous changes. Blue dots represent variants identified within duck samples, while red dots represent variants identified in human samples. Blank plots indicate that no variants were identified in that sample and gene.

**Figure S4: Neither diversity nor host-specific mutations increase over time**

(a) For each human sample, the full genome nucleotide diversity (π_Ν_ or π*_S_*) is plotted vs. the days post-symptom onset. Dark red dots represent the mean, full-genome nonsynonymous diversity for a given sample (π_Ν_), and light red dots represent the mean, full-genome synonymous diversity for that same sample (π*_S_*). Neither nonsynonymous nor synonymous diversity are correlated with days post symptom onset (nonsynonymous: r^2^ = −0.17, p = 0.69; synonymous: r^2^ = −0.22, p = −0.61). (b) To compare whether the number of putative host-adapting mutations increased over time in humans, we compared the number of host-specific and non-host specific mutations in humans sampled either in “early infection” (5-8 days post symptom onset), or in “late infection” (9-12 days post symptom onset). We divided the data into these categories by splitting on the mean days post symptom onset for human samples, which was 8 days. We then compared the proportion of host-specific variants during early and late infections with a Fisher’s exact test. The proportion of variants that are host-specific is not different in early vs. late infections (p = 0.72).

**Table S1: All within-host SNVs with annotations**

Every SNV identified in humans and ducks within-host are displayed with their frequency, coding region change, and functional annotation. All annotations for H5 HAs, N1 NAs, and all subtypes for all other genes were downloaded from the Influenza Research Database Sequence Feature Variant Types tool. Each SNV was then annotated as shown in the “description” column. These descriptions are paraphrased from annotations presented in the Influenza Research Database. We then manually curated annotated mutations to determine whether they were involved in “host-specific” functions or not, as shown in the “host-specific?” column. We defined host-specific functions/interactions as receptor binding, interaction with host cellular machinery, nuclear import and export, immune antagonism, 5’ cap binding, temperature sensitivity, and glycosylation. We also included sites that have been phenotypically identified as determinants of transmissibility and virulence. Sites that participate in binding interactions with other viral subunits or vRNP, conserved active site domains, drug resistance mutations, and epitope sites were not categorized as host-specific for this analysis. We annotated both synonymous and nonsynonymous mutations in our dataset.

