## Supplementary material for "Quantifying within-host diversity of H5N1 influenza viruses in humans and poultry in Cambodia": combined supplemental figures

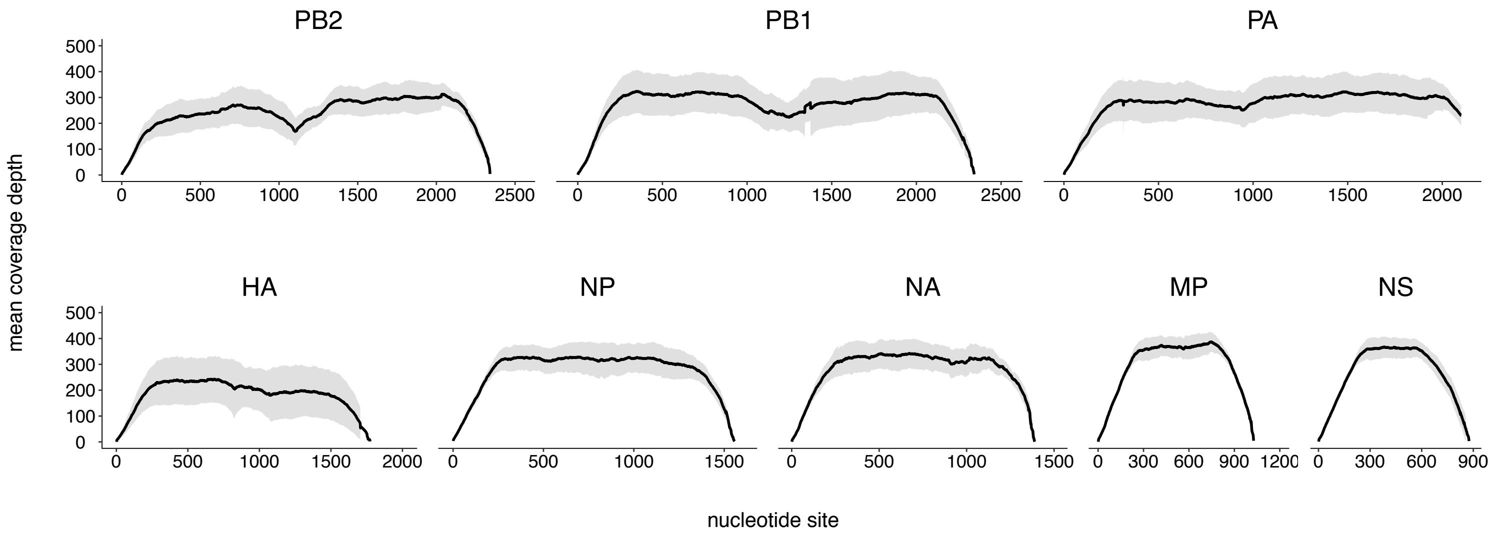


**Figure S1: Genome coverage**

The mean coverage depth at each nucleotide site (x-axis) for each gene across our 8 human and 5 duck samples is shown. Solid black lines represent the mean coverage across samples, and the grey shaded area represents the standard deviation of coverage depth across samples.

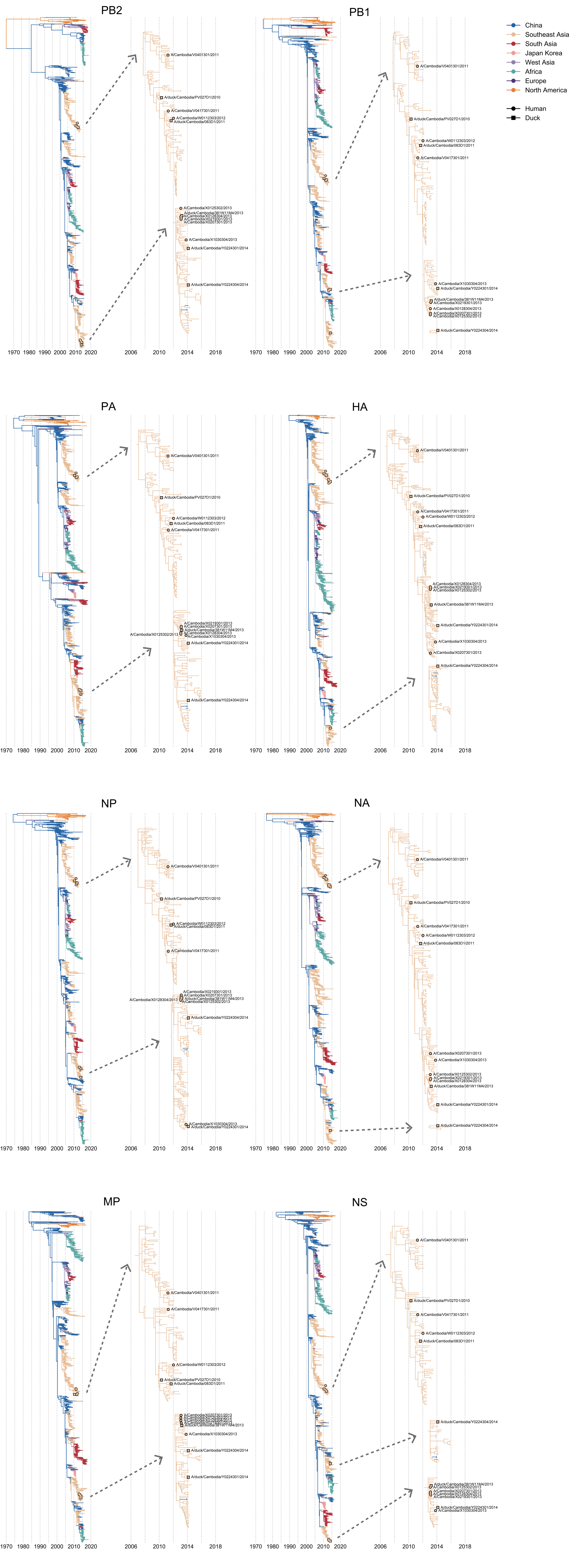


**Figure S2: Full genome phylogenetic placement of H5N1 virus samples from Cambodia**
All currently available H5N1 virus sequences were downloaded from the Influenza Research Database and the Global Initiative on Sharing All Influenza Data and used to generate full genome phylogenies using Nextstrain's augur pipeline. Colors represent the geographic region in which the sample was collected (for tips) or the inferred geographic location (for internal nodes). The x-axis position indicates the date of sample collection (for tips) or the inferred time to the most recent common ancestor (for internal nodes). In the full phylogenies (left), H5N1 viruses from Cambodia selected for within-host analysis are indicated by tan circles with black outlines. The subtrees containing the Cambodian samples selected for within-host analysis are shown to the right in the order that they appear in the full tree. In these trees, human tips are marked with a red circle, while duck tips are denoted with a blue circle. Internal genes from samples collected prior to 2013 belong to clade 1.1.2, while internal genes from samples collected in 2013 or later belong to clade 2.3.2.1a. All HA and NA sequences in this dataset, besides A/duck/Cambodia/Y0224304/2014, belong to clade 1.1.2.

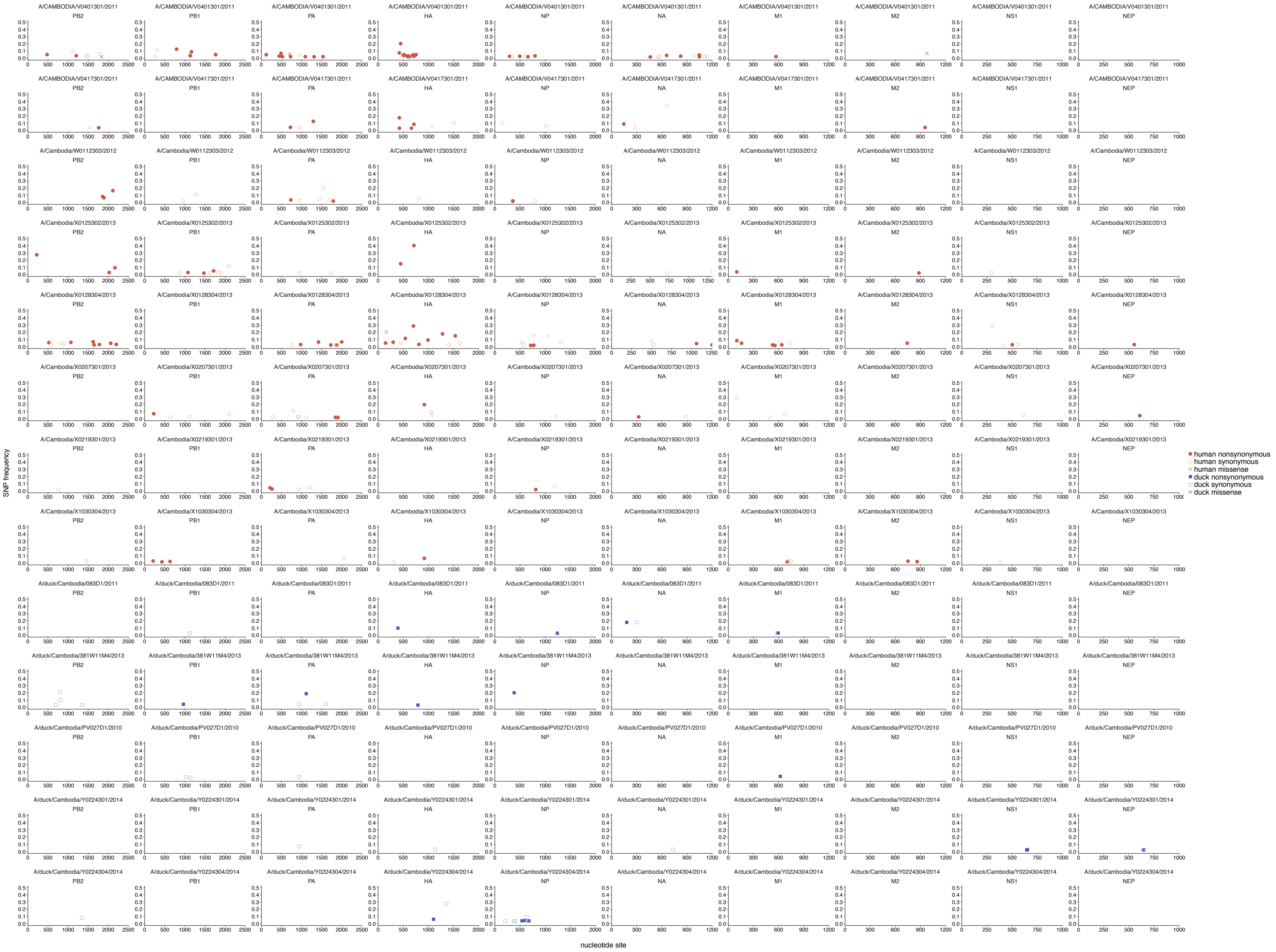

**Figure S3: All within-host variants detected in our dataset**
All within-host variants detected in our study are shown. Each row represents one sample and each column represents one gene. The x-axis shows the nucleotide site and the y-axis shows the frequency that the variant was detected within-host. Filled circles represent nonsynonymous changes, while open circles represent synonymous changes. Blue dots represent variants identified within duck samples, while red dots represent variants identified in human samples. Blank plots indicate that no variants were identified in that sample and gene.

**a**

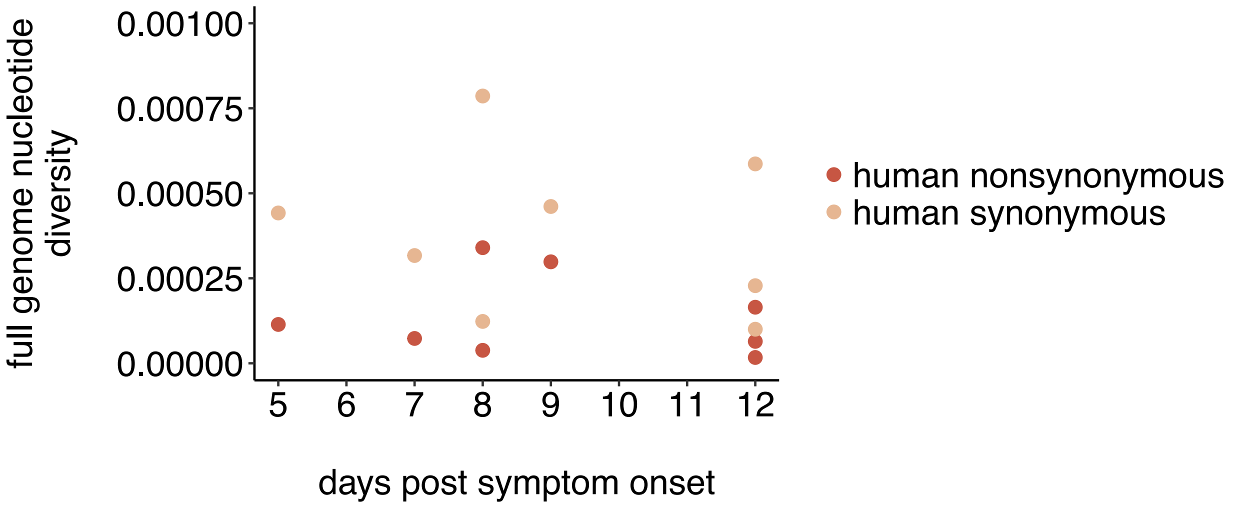


**b**

| | “early infection” (5-8 days) | “late infection” (9-12 days) |
|---|---|---|
| host-specific mutations | 18 | 22 |
| non-host specific mutations | 82 | 84 |

**Figure S4: Neither diversity nor host-specific mutations increase over time**
**(a)** For each human sample, the full genome nucleotide diversity ($\pi_N$ or $\pi_S$) is plotted vs. the days post-symptom onset. Dark red dots represent the mean, full-genome nonsynonymous diversity for a given sample ($\pi_N$), and light red dots represent the mean, full-genome synonymous diversity for that same sample ($\pi_S$). Neither nonsynonymous nor synonymous diversity are correlated with days post symptom onset (nonsynonymous: $r^2$ = -0.17, p = 0.69; synonymous: $r^2$ = -0.22, p = -0.61). **(b)** To compare whether the number of putative host-adapting mutations increased over time in humans, we compared the number of host-specific and non-host specific mutations in humans sampled either in “early infection” (5-8 days post symptom onset), or in “late infection” (9-12 days post symptom onset). We divided the data into these categories by splitting on the mean days post symptom onset for human samples, which was 8 days. We then compared the proportion of host-specific variants during early and late infections with a Fisher’s exact test. The proportion of variants that are host-specific is not different in early vs. late infections (p = 0.72).
