## supplemental table 1 for "Quantifying within-host diversity of H5N1 influenza viruses in humans and poultry in Cambodia"

**Table S1: All within-host SNVs with annotations**

| sample name | gene | site | ref | var | aa change | change type | freq (%) | description | Host-specific? |
| --- | --- | --- | --- | --- | --- | --- | --- | --- | --- |
| A/Cambodia/X0125302/2013 | PB2 | 220 | T | C | Met66Thr | nonsynonymous | 27.59% | No known function | no |
| A/Cambodia/V0401301/2011 | PB2 | 478 | A | C | His151Pro | nonsynonymous | 5.26% | No known function | no |
| A/Cambodia/X0128304/2013 | PB2 | 521 | A | T | Glu165Val | nonsynonymous | 5.88% | No known function | no |
| A/Cambodia/X0128304/2013 | PB2 | 573 | A | G | Gln182Gln | synonymous | 7.79% | No known function | no |
| A/Cambodia/X0128304/2013 | PB2 | 574 | T | C | Leu183Leu | synonymous | 2.61% | No known function | no |
| A/duck/Cambodia/381W11M4/2013 | PB2 | 693 | C | T | Gly222Gly | synonymous | 3.27% | No known function | no |
| A/Cambodia/X0219301/2013 | PB2 | 758 | T | G | Thr245Thr | synonymous | 2.18% | No known function | no |
| A/duck/Cambodia/381W11M4/2013 | PB2 | 798 | A | G | Gln257Gln | synonymous | 21.50% | No known function | no |
| A/duck/Cambodia/381W11M4/2013 | PB2 | 813 | T | C | Ala262Ala | synonymous | 10.28% | No known function | no |
| A/Cambodia/X0128304/2013 | PB2 | 819 | A | G | Arg264Arg | synonymous | 5.31% | No known function | no |
| A/Cambodia/X0128304/2013 | PB2 | 897 | T | C | Gly290Gly | synonymous | 4.39% | No known function | no |
| A/Cambodia/X0128304/2013 | PB2 | 1069 | A | T | Asn348Tyr | nonsynonymous | 6.15% | Putative m7GTP cap binding site. | yes |
| A/Cambodia/V0401301/2011 | PB2 | 1115 | C | T | Phe363Phe | synonymous | 10% | Putative m7GTP cap binding site. | yes |
| A/Cambodia/V0401301/2011 | PB2 | 1202 | A | C | Gln392His | nonsynonymous | 3.61% | Putative m7GTP cap binding site. | yes |
| A/duck/Cambodia/Y0224304/2014 | PB2 | 1340 | C | T | Asp441Asp | synonymous | 8.16% | Putative m7GTP cap binding site. | yes |
| A/duck/Cambodia/381W11M4/2013 | PB2 | 1350 | C | T | Asp441Asp | synonymous | 3.19% | Putative m7GTP cap binding site. | yes |
| A/Cambodia/X1030304/2013 | PB2 | 1451 | A | G | Arg479Arg | synonymous | 2.87% | Nuclear localization motif. | yes |
| A/Cambodia/V0401301/2011 | PB2 | 1460 | A | G | Val478Val | synonymous | 3.90% | Nuclear localization motif. | yes |
| A/Cambodia/V0401301/2011 | PB2 | 1484 | T | C | Asp486Asp | synonymous | 2.30% | Nuclear localization motif. | yes |
| A/Cambodia/V0417301/2011 | PB2 | 1539 | C | A | Arg505Arg | synonymous | 4.05% | No known function | no |
| A/Cambodia/X0128304/2013 | PB2 | 1621 | T | C | Ser532Pro | nonsynonymous | 6.90% | No known function | no |
| A/Cambodia/X0128304/2013 | PB2 | 1646 | A | G | Asn540Ser | nonsynonymous | 2.65% | No known function | no |
| A/Cambodia/V0417301/2011 | PB2 | 1761 | C | T | Pro579Ser | nonsynonymous | 3.80% | No known function | no |
| A/Cambodia/X0128304/2013 | PB2 | 1778 | T | C | Val584Ala | nonsynonymous | 3.01% | No known function | no |
| A/Cambodia/V0401301/2011 | PB2 | 1784 | A | G | Lys586Lys | synonymous | 6.01% | No known function | no |
| A/Cambodia/V0401301/2011 | PB2 | 1830 | C | T | Gln602Stop | stop gained | 2.09% | No known function | no |
| A/Cambodia/W0112303/2012 | PB2 | 1859 | T | C | Ile616Thr | nonsynonymous | 8.43% | No known function | no |
| A/Cambodia/W0112303/2012 | PB2 | 1891 | G | A | Glu627Lys | nonsynonymous | 6.63% | A Lys at 627 enhances mammalian replication. | yes |
| A/Cambodia/X0125302/2013 | PB2 | 2022 | G | A | Val667Ile | nonsynonymous | 2.99% | An Ile at 667 was associated with human-infecting H5N1 strain. | yes |
| A/Cambodia/X0128304/2013 | PB2 | 2060 | A | T | Asp678Val | nonsynonymous | 4.80% | No known function | no |
| A/Cambodia/W0112303/2012 | PB2 | 2113 | A | G | Asn701Asp | nonsynonymous | 16.49% | An Asn at 701 enhances mammalian replication. | yes |
| A/Cambodia/X0125302/2013 | PB2 | 2163 | A | G | Ser714Gly | nonsynonymous | 9.59% | An Arg at 714 enhances mammalian replication. | yes |
| A/Cambodia/X0128304/2013 | PB2 | 2198 | T | A | Val724Glu | nonsynonymous | 3.41% | No known function | no |
| A/Cambodia/X1030304/2013 | PB1 | 212 | G | A | Gly71Glu | nonsynonymous | 3.12% | No known function | no |
| A/Cambodia/X0207301/2013 | PB1 | 226 | G | A | Asp76Asn | nonsynonymous | 7.14% | No known function | no |
| A/Cambodia/V0401301/2011 | PB1 | 255 | A | G | Thr85Thr | synonymous | 2.12% | No known function | no |
| A/Cambodia/V0401301/2011 | PB1 | 312 | A | G | Glu104Glu | synonymous | 11.42% | No known function | no |
| A/Cambodia/X1030304/2013 | PB1 | 431 | C | T | Ala144Val | nonsynonymous | 1.78% | No known function | no |
| A/Cambodia/X1030304/2013 | PB1 | 631 | A | G | Arg211Gly | nonsynonymous | 2.34% | Nuclear localization motif. | yes |
| A/Cambodia/X0207301/2013 | PB1 | 634 | C | T | Leu212Leu | synonymous | 2.10% | Nuclear localization motif. | yes |
| A/Cambodia/V0401301/2011 | PB1 | 794 | A | G | Lys265Arg | nonsynonymous | 12.65% | No known function | no |
| A/Cambodia/X0125302/2013 | PB1 | 857 | A | G | Lys279Lys | synonymous | 2.14% | No known function | no |
| A/Cambodia/W0112303/2012 | PB1 | 963 | G | A | Gln313Gln | synonymous | 2.10% | No known function | no |
| A/duck/Cambodia/381W11M4/2013 | PB1 | 968 | A | G | Met317Val | nonsynonymous | 4.21% | No known function | no |
| A/duck/Cambodia/PV027D1/2010 | PB1 | 1026 | G | A | Arg334Arg | synonymous | 3.30% | No known function | no |
| A/Cambodia/X0125302/2013 | PB1 | 1078 | A | G | Lys353Arg | nonsynonymous | 2.94% | An Arg at 353 is associated with higher replication and pathogenicity of an H1N1 pandemic strain. | yes |
| A/Cambodia/V0401301/2011 | PB1 | 1113 | A | G | Glu371Glu | synonymous | 4.33% | No known function | no |
| A/Cambodia/X0207301/2013 | PB1 | 1113 | A | G | Glu371Glu | synonymous | 3.15% | No known function | no |
| A/duck/Cambodia/083D1/2011 | PB1 | 1121 | A | G | Glu371Glu | synonymous | 2.71% | No known function | no |
| A/Cambodia/V0401301/2011 | PB1 | 1136 | A | G | Lys379Arg | nonsynonymous | 3.64% | No known function | no |
| A/duck/Cambodia/PV027D1/2010 | PB1 | 1137 | A | G | Glu371Glu | synonymous | 2.59% | No known function | no |
| A/Cambodia/V0401301/2011 | PB1 | 1165 | A | G | Ile389Val | nonsynonymous | 8.87% | No known function | no |
| A/Cambodia/W0112303/2012 | PB1 | 1267 | C | T | Leu415Leu | synonymous | 11.21% | No known function | no |
| A/Cambodia/X0125302/2013 | PB1 | 1472 | A | G | Ile484Met | nonsynonymous | 2.17% | No known function | no |
| A/Cambodia/X0125302/2013 | PB1 | 1631 | C | T | Asn537Asn | synonymous | 2.50% | No known function | no |
| A/Cambodia/X0125302/2013 | PB1 | 1716 | A | T | Thr566Ser | nonsynonymous | 5.20% | An Ala at 566 is associated with higher replication and pathogenicity of an H1N1 pandemic virulence. | yes |
| A/Cambodia/V0401301/2011 | PB1 | 1758 | A | G | Lys586Lys | synonymous | 5.70% | No known function | no |
| A/Cambodia/V0401301/2011 | PB1 | 1766 | T | C | Leu589Pro | nonsynonymous | 5% | No known function | no |
| A/Cambodia/X0125302/2013 | PB1 | 1823 | C | T | Ile601Ile | synonymous | 2.75% | No known function | no |
| A/Cambodia/X0125302/2013 | PB1 | 1901 | T | C | Pro627Pro | synonymous | 3.08% | No known function | no |
| A/Cambodia/X0125302/2013 | PB1 | 2090 | A | G | Gln690Gln | synonymous | 11.89% | 690 falls into a region of PB1 that binds to PB2 and is critical for PB2 enzymatic activity. | no |
| A/Cambodia/X0207301/2013 | PB1 | 2100 | T | C | Phe700Phe | synonymous | 7.20% | 700 falls into a region of PB1 that binds to PB2 and is critical for PB2 enzymatic activity. | no |
| A/Cambodia/V0401301/2011 | PA | 122 | T | C | Phe35Ser | nonsynonymous | 5% | No known function | no |
| A/Cambodia/X0219301/2013 | PA | 215 | C | T | Pro68Leu | nonsynonymous | 4.32% | No known function | no |
| A/Cambodia/X0219301/2013 | PA | 265 | A | G | Thr85Ala | nonsynonymous | 2.84% | An Ile at 85 enhances polymerase activity of pandemic H1N1 in mammalian cell. | yes |
| A/Cambodia/X0207301/2013 | PA | 291 | T | C | Ser93Ser | synonymous | 2.22% | No known function | no |
| A/Cambodia/V0401301/2011 | PA | 443 | G | A | Arg142Lys | nonsynonymous | 3.07% | No known function | no |
| A/Cambodia/V0401301/2011 | PA | 488 | C | A | Thr157Asn | nonsynonymous | 6.85% | No known function | no |
| A/Cambodia/V0401301/2011 | PA | 523 | G | A | Ala169Thr | nonsynonymous | 2.24% | No known function | no |
| A/Cambodia/V0401301/2011 | PA | 706 | A | T | Arg230Stop | stop gained | 5.94% | No known function | no |
| A/Cambodia/V0417301/2011 | PA | 723 | A | G | Lys237Glu | nonsynonymous | 4.36% | No known function | no |
| A/Cambodia/V0401301/2011 | PA | 727 | A | G | Lys237Glu | nonsynonymous | 2.34% | No known function | no |
| A/Cambodia/W0112303/2012 | PA | 732 | A | G | Lys237Glu | nonsynonymous | 3.64% | No known function | no |
| A/Cambodia/X0128304/2013 | PA | 765 | T | C | Ser247Ser | synonymous | 2.24% | No known function | no |
| A/Cambodia/X0207301/2013 | PA | 789 | A | G | Pro259Pro | synonymous | 11.21% | This site falls into a region of PA that interacts with PB1 residues 1-25, and this interaction is critical for transcription. | no |
| A/Cambodia/X0207301/2013 | PA | 906 | G | A | Glu298Glu | synonymous | 2.26% | This site falls into a region of PA that interacts with PB1 residues 1-25, and this interaction is critical for transcription. | no |
| A/Cambodia/X0207301/2013 | PA | 933 | A | G | Ala307Ala | synonymous | 2.80% | This site falls into a region of PA that interacts with PB1 residues 1-25, and this interaction is critical for transcription. | no |
| A/Cambodia/X0219301/2013 | PA | 933 | A | G | Ala307Ala | synonymous | 2.37% | This site falls into a region of PA that interacts with PB1 residues 1-25, and this interaction is critical for transcription. | no |
| A/Cambodia/V0417301/2011 | PA | 935 | A | G | Ala307Ala | synonymous | 4.38% | This site falls into a region of PA that interacts with PB1 residues 1-25, and this interaction is critical for transcription. | no |
| A/Cambodia/V0401301/2011 | PA | 939 | A | G | Ala307Ala | synonymous | 4.67% | This site falls into a region of PA that interacts with PB1 residues 1-25, and this interaction is critical for transcription. | no |
| A/Cambodia/X0125302/2013 | PA | 939 | A | G | Ala307Ala | synonymous | 2.50% | This site falls into a region of PA that interacts with PB1 residues 1-25, and this interaction is critical for transcription. | no |
| A/duck/Cambodia/381W11M4/2013 | PA | 939 | A | G | Ala307Ala | synonymous | 4.55% | This site falls into a region of PA that interacts with PB1 residues 1-25, and this interaction is critical for transcription. | no |
| A/duck/Cambodia/PV027D1/2010 | PA | 941 | A | G | Ala307Ala | synonymous | 3.31% | This site falls into a region of PA that interacts with PB1 residues 1-25, and this interaction is critical for transcription. | no |
| A/Cambodia/W0112303/2012 | PA | 944 | A | G | Ala307Ala | synonymous | 2.85% | This site falls into a region of PA that interacts with PB1 residues 1-25, and this interaction is critical for transcription. | no |
| A/duck/Cambodia/Y0224301/2014 | PA | 945 | A | G | Ala307Ala | synonymous | 7.28% | This site falls into a region of PA that interacts with PB1 residues 1-25, and this interaction is critical for transcription. | no |
| A/Cambodia/X0128304/2013 | PA | 976 | A | G | Lys318Glu | nonsynonymous | 3.05% | This site falls into a region of PA that interacts with PB1 residues 1-25, and this interaction is critical for transcription. | no |
| A/Cambodia/V0401301/2011 | PA | 1094 | A | C | Asn359Thr | nonsynonymous | 2.26% | This site falls into a region of PA that interacts with PB1 residues 1-25, and this interaction is critical for transcription. | no |
| A/Cambodia/X0207301/2013 | PA | 1110 | G | A | Leu366Leu | synonymous | 2.08% | This site falls into a region of PA that interacts with PB1 residues 1-25, and this interaction is critical for transcription. | no |
| A/duck/Cambodia/381W11M4/2013 | PA | 1118 | G | A | Arg367Lys | nonsynonymous | 19% | This site falls into a region of PA that interacts with PB1 residues 1-25, and this interaction is critical for transcription. | no |
| A/Cambodia/X0219301/2013 | PA | 1209 | G | A | Lys399Lys | synonymous | 4.79% | This site falls into a region of PA that interacts with PB1 residues 1-25, and this interaction is critical for transcription. | no |
| A/Cambodia/V0417301/2011 | PA | 1292 | T | A | Asp426Glu | nonsynonymous | 12.60% | This site falls into a region of PA that interacts with PB1 residues 1-25, and this interaction is critical for transcription. | no |
| A/Cambodia/V0401301/2011 | PA | 1312 | G | A | Val432Ile | nonsynonymous | 2.08% | This site falls into a region of PA that interacts with PB1 residues 1-25, and this interaction is critical for transcription. | no |
| A/Cambodia/X0128304/2013 | PA | 1420 | A | C | Asn466His | nonsynonymous | 6.45% | This site falls into a region of PA that interacts with PB1 residues 1-25, and this interaction is critical for transcription. | no |
| A/Cambodia/W0112303/2012 | PA | 1428 | C | T | Leu469Leu | synonymous | 4.59% | This site falls into a region of PA that interacts with PB1 residues 1-25, and this interaction is critical for transcription. | no |
| A/Cambodia/V0401301/2011 | PA | 1533 | A | G | Ile505Met | nonsynonymous | 2.24% | This site falls into a region of PA that interacts with PB1 residues 1-25, and this interaction is critical for transcription. | no |
| A/Cambodia/W0112303/2012 | PA | 1544 | A | G | Gly507Gly | synonymous | 20.26% | This site falls into a region of PA that interacts with PB1 residues 1-25, and this interaction is critical for transcription. | no |
| A/duck/Cambodia/381W11M4/2013 | PA | 1608 | G | A | Pro530Pro | synonymous | 4.38% | This site falls into a region of PA that interacts with PB1 residues 1-25, and this interaction is critical for transcription. | no |
| A/Cambodia/W0112303/2012 | PA | 1691 | A | G | Gln556Gln | synonymous | 3.60% | This site falls into a region of PA that interacts with PB1 residues 1-25, and this interaction is critical for transcription. | no |
| A/Cambodia/X0128304/2013 | PA | 1727 | A | G | Asn568Ser | nonsynonymous | 2.40% | This site falls into a region of PA that interacts with PB1 residues 1-25, and this interaction is critical for transcription. | no |
| A/Cambodia/X0125302/2013 | PA | 1728 | G | A | Thr570Thr | synonymous | 2.03% | This site falls into a region of PA that interacts with PB1 residues 1-25, and this interaction is critical for transcription. | no |
| A/Cambodia/W0112303/2012 | PA | 1789 | T | C | Leu589Pro | nonsynonymous | 2.07% | This site falls into a region of PA that interacts with PB1 residues 1-25, and this interaction is critical for transcription. | no |
| A/Cambodia/X0207301/2013 | PA | 1841 | A | G | Glu610Gly | nonsynonymous | 2.15% | This site falls into a region of PA that interacts with PB1 residues 1-25, and this interaction is critical for transcription. | no |
| A/Cambodia/X0128304/2013 | PA | 1868 | A | G | Lys615Arg | nonsynonymous | 2.47% | An Asn at PA 615 has been associated with adaptation of avian influenza polymerases to humans. | yes |
| A/Cambodia/X0128304/2013 | PA | 1893 | A | G | Glu623Glu | synonymous | 2.52% | This site falls into a region of PA that interacts with PB1 residues 1-25, and this interaction is critical for transcription. | no |
| A/Cambodia/X0207301/2013 | PA | 1902 | A | G | Glu630Glu | synonymous | 3.07% | This site falls into a region of PA that interacts with PB1 residues 1-25, and this interaction is critical for transcription. | no |
| A/Cambodia/X0207301/2013 | PA | 1903 | A | G | Ser631Gly | nonsynonymous | 1.79% | A Ser at 631 enhances virulence of H5N1 in mice. | yes |
| A/Cambodia/X0128304/2013 | PA | 1999 | T | C | Ser659Pro | nonsynonymous | 6.79% | This site falls into a region of PA that interacts with PB1 residues 1-25, and this interaction is critical for transcription. | no |
| A/Cambodia/X1030304/2013 | PA | 2055 | A | G | Gly679Gly | synonymous | 6.98% | This site falls into a region of PA that interacts with PB1 residues 1-25, and this interaction is critical for transcription. | no |
| A/Cambodia/X0128304/2013 | HA | 149 | C | T | Thr41Ile | Nonsynonymous | 5.19% | No known function. | no |
| A/Cambodia/X0128304/2013 | HA | 163 | C | T | Gln46Stop | stop gained | 20.24% | No known function. | no |
| A/Cambodia/X0128304/2013 | HA | 299 | A | G | Glu91Gly | nonsynonymous | 6.33% | A Lys at 91 enhances α-2,6 binding. | yes |
| A/Cambodia/X1030304/2013 | HA | 306 | C | T | Val102Val | synonymous | 1.79% | No known function. | no |
| A/duck/Cambodia/083D1/2011 | HA | 394 | A | G | Lys129Glu | nonsynonymous | 9.63% | No known function. | no |
| A/Cambodia/V0401301/2011 | HA | 422 | A | G | His141Arg | nonsynonymous | 7.56% | No known function. | no |
| A/Cambodia/V0417301/2011 | HA | 422 | A | G | His141Arg | nonsynonymous | 17.50% | No known function. | no |
| A/Cambodia/V0417301/2011 | HA | 425 | A | G | Glu142Gly | nonsynonymous | 3.20% | Putative glycosylation site. | yes |
| A/Cambodia/V0401301/2011 | HA | 449 | C | T | Ala150Val | nonsynonymous | 20.24% | A Val at 150 confers enhanced α-2,6 sialic acid binding in H5N1 viruses. | yes |
| A/Cambodia/X0125302/2013 | HA | 449 | C | T | Ala150Val | nonsynonymous | 15.09% | A Val at 150 confers enhanced α-2,6 sialic acid binding in H5N1 viruses. | yes |
| A/Cambodia/X0128304/2013 | HA | 450 | T | C | His141His | synonymous | 5.69% | No known function | no |
| A/Cambodia/V0401301/2011 | HA | 497 | T | C | Leu166Pro | nonsynonymous | 4.28% | No known function | no |
| A/Cambodia/V0401301/2011 | HA | 513 | T | C | Ser171Ser | synonymous | 3.30% | Part of putative glycosylation motif that improves α-2,6 binding. | yes |
| A/Cambodia/V0401301/2011 | HA | 517 | T | C | Tyr173His | nonsynonymous | 5.04% | Residue involved in sialic acid recognition. | yes |
| A/Cambodia/V0401301/2011 | HA | 527 | T | C | Ile176Thr | nonsynonymous | 4.26% | No known function | no |
| A/Cambodia/X0128304/2013 | HA | 542 | A | C | Lys172Thr | nonsynonymous | 11.50% | Part of putative glycosylation motif that improves α-2,6 binding. | yes |
| A/Cambodia/V0401301/2011 | HA | 590 | C | T | Pro197Leu | nonsynonymous | 2.40% | No known function | no |
| A/Cambodia/V0401301/2011 | HA | 593 | A | G | Asn198Ser | nonsynonymous | 3.32% | A Lys at 198 confers α-2,6 sialic acid binding. | yes |
| A/Cambodia/V0401301/2011 | HA | 628 | C | A | Pro210Thr | nonsynonymous | 2.42% | No known function | no |
| A/Cambodia/V0417301/2011 | HA | 664 | A | G | Asn222Asp | nonsynonymous | 3.18% | No known function | no |
| A/Cambodia/V0401301/2011 | HA | 694 | A | G | Arg232Gly | nonsynonymous | 4.83% | No known function | no |
| A/Cambodia/V0401301/2011 | HA | 695 | G | A | Arg232Lys | nonsynonymous | 4.07% | No known function | no |
| A/Cambodia/X0128304/2013 | HA | 703 | A | G | Thr226Ala | nonsynonymous | 28.91% | An Ile at 226 enhanced α-2,6 sialic acid binding | yes |
| A/Cambodia/V0401301/2011 | HA | 713 | A | T | Gln238Leu | nonsynonymous | 2.80% | A Leu at 238 confers a switch from α-2,3 to α-2,6 sialic acid binding and is a determinant of mammalian transmission. | yes |
| A/Cambodia/V0417301/2011 | HA | 713 | A | T | Gln238Leu | nonsynonymous | 8.45% | A Leu at 238 confers a switch from α-2,3 to α-2,6 sialic acid binding and is a determinant of mammalian transmission. | yes |
| A/Cambodia/X0125302/2013 | HA | 713 | A | G | Gln238Arg | nonsynonymous | 40.30% | A Leu at 238 confers a switch from α-2,3 to α-2,6 sialic acid binding and is a determinant of mammalian transmission. | yes |
| A/Cambodia/V0401301/2011 | HA | 754 | A | G | Asn252Asp | nonsynonymous | 5.08% | No known function | no |
| A/duck/Cambodia/381W11M4/2013 | HA | 793 | G | A | Ala265Thr | nonsynonymous | 3.28% | No known function | no |
| A/Cambodia/W0112303/2012 | HA | 806 | T | C | Ala265Ala | synonymous | 5.62% | No known function | no |
| A/Cambodia/X0128304/2013 | HA | 811 | A | G | Asn262Asp | nonsynonymous | 3.38% | No known function | no |
| A/Cambodia/X0207301/2013 | HA | 919 | T | C | Phe307Leu | nonsynonymous | 19.82% | No known function | no |
| A/Cambodia/X1030304/2013 | HA | 919 | T | C | Phe307Leu | nonsynonymous | 6.65% | No known function | no |
| A/Cambodia/X0128304/2013 | HA | 992 | T | C | Val322Ala | nonsynonymous | 9.30% | No known function | no |
| A/Cambodia/X0207301/2013 | HA | 1056 | A | C | Ile352Ile | synonymous | 9.42% | No known function | no |
| A/Cambodia/V0417301/2011 | HA | 1071 | A | G | Glu357Glu | synonymous | 6.35% | No known function | no |
| A/Cambodia/X0207301/2013 | HA | 1071 | A | G | Glu357Glu | synonymous | 5.83% | No known function | no |
| A/duck/Cambodia/Y0224304/2014 | HA | 1103 | G | A | Val363Ile | nonsynonymous | 6.32% | No known function | no |
| A/duck/Cambodia/Y0224301/2014 | HA | 1131 | A | T | Gly377Gly | synonymous | 3% | No known function | no |
| A/Cambodia/X0219301/2013 | HA | 1221 | C | T | Thr407Thr | synonymous | 17.86% | No known function | no |
| A/Cambodia/X0128304/2013 | HA | 1281 | C | A | Asn418Lys | nonsynonymous | 27.90% | No known function | no |
| A/duck/Cambodia/Y0224304/2014 | HA | 1360 | A | G | Glu448Glu | synonymous | 2.37% | No known function | no |
| A/Cambodia/X0128304/2013 | HA | 1410 | A | G | Val461Val | synonymous | 10.74% | No known function | no |
| A/Cambodia/V0417301/2011 | HA | 1506 | A | G | Thr502Thr | synonymous | 15.22% | No known function | no |
| A/Cambodia/X0128304/2013 | HA | 1535 | A | G | Tyr503Cys | nonsynonymous | 4.98% | No known function | no |
| A/Cambodia/X0128304/2013 | HA | 1629 | A | G | Ser534Ser | synonymous | 10.14% | No known function | no |
| A/Cambodia/V0417301/2011 | NP | 146 | G | A | Gly37Gly | synonymous | 3.90% | RNA-binding domain | no |
| A/duck/Cambodia/Y0224304/2014 | NP | 207 | C | T | Asn59Asn | synonymous | 2.70% | RNA-binding domain | no |
| A/Cambodia/V0401301/2011 | NP | 293 | G | A | Arg98Gln | nonsynonymous | 2.42% | RNA-binding domain | no |
| A/Cambodia/V0401301/2011 | NP | 342 | G | A | Glu114Glu | synonymous | 1.96% | RNA-binding domain | no |
| A/Cambodia/W0112303/2012 | NP | 356 | T | C | Leu108Pro | nonsynonymous | 2.23% | RNA-binding domain | no |
| A/Cambodia/W0112303/2012 | NP | 358 | A | G | Ile109Val | nonsynonymous | 3.20% | RNA-binding domain | no |
| A/duck/Cambodia/Y0224304/2014 | NP | 378 | C | T | Ile116Ile | synonymous | 20.43% | RNA-binding domain | no |
| A/duck/Cambodia/381W11M4/2013 | NP | 384 | A | G | Gln117Arg | nonsynonymous | 3.53% | RNA-binding domain | no |
| A/duck/Cambodia/Y0224304/2014 | NP | 402 | C | T | Asn124Asn | synonymous | 2.87% | RNA-binding domain | no |
| A/Cambodia/V0401301/2011 | NP | 496 | C | G | Leu166Val | nonsynonymous | 3.77% | RNA-binding domain | no |
| A/duck/Cambodia/Y0224304/2014 | NP | 539 | C | T | Ser170Leu | nonsynonymous | 6.13% | RNA-binding domain | no |
| A/Cambodia/X0128304/2013 | NP | 542 | A | G | Gln168Gln | synonymous | 1.92% | RNA-binding domain | no |
| A/duck/Cambodia/Y0224304/2014 | NP | 593 | C | T | Thr188Ile | nonsynonymous | 4.76% | No known function | no |
| A/Cambodia/V0401301/2011 | NP | 603 | C | A | Val201Val | synonymous | 2.81% | Nuclear targeting motif | yes |
| A/duck/Cambodia/Y0224304/2014 | NP | 633 | C | T | Ile201Ile | synonymous | 9.23% | Nuclear targeting motif | yes |
| A/duck/Cambodia/Y0224304/2014 | NP | 636 | C | T | Asn202Asn | synonymous | 3.70% | Nuclear targeting motif | yes |
| A/Cambodia/V0401301/2011 | NP | 659 | A | G | Glu220Gly | nonsynonymous | 2.17% | Region involved in NP-NP association | no |
| A/duck/Cambodia/Y0224304/2014 | NP | 674 | C | T | Thr215Ile | nonsynonymous | 3.69% | Nuclear targeting motif | yes |
| A/Cambodia/X0128304/2013 | NP | 712 | T | C | Ile225Thr | nonsynonymous | 1.82% | Region involved in NP-NP association | no |
| A/Cambodia/X0128304/2013 | NP | 770 | G | A | Glu244Glu | synonymous | 15.70% | Region involved in NP-NP association | no |
| A/Cambodia/X0128304/2013 | NP | 771 | A | G | Ser245Gly | nonsynonymous | 2.04% | Region involved in NP-NP association | no |
| A/Cambodia/X0128304/2013 | NP | 774 | A | G | Arg246Gly | nonsynonymous | 2.24% | Region involved in NP-NP association | no |
| A/Cambodia/V0401301/2011 | NP | 799 | A | G | Arg267Gly | nonsynonymous | 3.36% | Region involved in NP-NP association | no |
| A/Cambodia/W0112303/2012 | NP | 804 | T | A | Ile257Ile | synonymous | 1.92% | Region involved in NP-NP association | no |
| A/Cambodia/X0219301/2013 | NP | 814 | C | T | Ala260Val | nonsynonymous | 2.02% | Region involved in NP-NP association | no |
| A/Cambodia/V0417301/2011 | NP | 1017 | T | C | Leu328Leu | synonymous | 6.72% | Region involved in NP-NP association | no |
| A/Cambodia/X0128304/2013 | NP | 1055 | G | A | Glu339Glu | synonymous | 14.89% | Region involved in NP-NP association | no |
| A/Cambodia/X0219301/2013 | NP | 1184 | T | C | Ser383Ser | synonymous | 6.30% | Region involved in NP-NP association | no |
| A/Cambodia/X0207301/2013 | NP | 1217 | G | A | Arg391Arg | synonymous | 3.81% | Region involved in NP-NP association | no |
| A/duck/Cambodia/083D1/2011 | NP | 1241 | C | T | Ala403Val | nonsynonymous | 2.92% | Region involved in NP-NP association | no |
| A/duck/Cambodia/083D1/2011 | NP | 1242 | A | T | Ala403Ala | synonymous | 2.52% | Region involved in NP-NP association | no |
| A/Cambodia/X0128304/2013 | NP | 1319 | A | G | Ala427Ala | synonymous | 6.31% | Region involved in NP-NP association | no |
| A/Cambodia/V0417301/2011 | NA | 146 | G | A | Glu47Lys | nonsynonymous | 8.89% | A deletion in this part of NA has been linked to host range. | yes |
| A/duck/Cambodia/083D1/2011 | NA | 181 | A | G | Lys58Glu | nonsynonymous | 17.89% | A deletion in this part of NA has been linked to host range. | yes |
| A/Cambodia/V0417301/2011 | NA | 280 | A | G | Lys91Lys | synonymous | 3.70% | No known function | no |
| A/duck/Cambodia/083D1/2011 | NA | 297 | C | A | Val96Val | synonymous | 17.95% | No known function | no |
| A/Cambodia/X0207301/2013 | NA | 323 | C | A | His106Gln | nonsynonymous | 2.83% | No known function | no |
| A/Cambodia/V0401301/2011 | NA | 462 | C | G | Pro149Arg | nonsynonymous | 1.92% | An Ala at 149 is responsible for reduced zanamivir sensitivity in an H5N1 virus. 149 is part of the NA catalytic site. | no |
| A/Cambodia/X0128304/2013 | NA | 497 | C | T | Cys164Cys | synonymous | 8.01% | No known function | no |
| A/Cambodia/X0128304/2013 | NA | 527 | A | G | Gly174Gly | synonymous | 2.95% | No known function | no |
| A/Cambodia/V0401301/2011 | NA | 553 | C | T | Asp179Asp | synonymous | 1.73% | Oseltamivir binding site | no |
| A/Cambodia/V0401301/2011 | NA | 571 | A | G | Val185Val | synonymous | 2.49% | No known function | no |
| A/Cambodia/V0401301/2011 | NA | 656 | G | A | Val214Ile | nonsynonymous | 3.98% | No known function | no |
| A/Cambodia/V0417301/2011 | NA | 658 | T | C | Tyr217Tyr | synonymous | 34.25% | No known function | no |
| A/Cambodia/X0125302/2013 | NA | 722 | A | G | Glu239Glu | synonymous | 1.82% | No known function | no |
| A/duck/Cambodia/Y0224301/2014 | NA | 733 | A | G | Lys242Lys | synonymous | 2.65% | No known function | no |
| A/Cambodia/V0401301/2011 | NA | 824 | T | A | Cys270Ser | nonsynonymous | 3.01% | No known function | no |
| A/Cambodia/X0207301/2013 | NA | 881 | T | C | Tyr292Tyr | synonymous | 3.15% | No known function | no |
| A/Cambodia/V0401301/2011 | NA | 985 | A | G | Ala323Ala | synonymous | 3.80% | No known function | no |
| A/Cambodia/V0401301/2011 | NA | 1046 | A | G | Ser344Gly | nonsynonymous | 4.56% | Oseltamivir binding site | no |
| A/Cambodia/V0401301/2011 | NA | 1047 | G | A | Ser344Asn | nonsynonymous | 2.11% | Oseltamivir binding site | no |
| A/Cambodia/X0128304/2013 | NA | 1062 | A | G | Met353Val | nonsynonymous | 4.51% | No known function | no |
| A/Cambodia/V0401301/2011 | NA | 1108 | C | T | Asp364Asp | synonymous | 2.42% | No known function | no |
| A/Cambodia/V0401301/2011 | NA | 1138 | A | G | Val374Val | synonymous | 3.09% | No known function | no |
| A/Cambodia/X0128304/2013 | NA | 1257 | A | G | Thr418Ala | nonsynonymous | 2.52% | No known function | no |
| A/Cambodia/X0125302/2013 | NA | 1304 | G | A | Val433Val | synonymous | 4.63% | No known function | no |
| A/Cambodia/X0207301/2013 | M1 | 100 | G | A | Lys27Lys | synonymous | 29.01% | Membrane-binding region | no |
| A/Cambodia/X0125302/2013 | M1 | 101 | G | A | Asp30Asn | nonsynonymous | 3.93% | Membrane-binding region | no |
| A/Cambodia/X0128304/2013 | M1 | 102 | A | G | Gln26Arg | nonsynonymous | 8.57% | Membrane-binding region | no |
| A/Cambodia/X0128304/2013 | M1 | 158 | T | G | Trp45Gly | nonsynonymous | 5.10% | Membrane-binding region | no |
| A/Cambodia/X0207301/2013 | M1 | 496 | T | C | His159His | synonymous | 1.96% | Membrane-binding region | no |
| A/Cambodia/X0128304/2013 | M1 | 528 | T | C | Ile168Thr | nonsynonymous | 2.61% | vRNP binding region | no |
| A/Cambodia/X0128304/2013 | M1 | 557 | A | G | Arg178Gly | nonsynonymous | 1.80% | vRNP binding region | no |
| A/Cambodia/V0401301/2011 | M1 | 569 | G | A | Gly185Asp | nonsynonymous | 2.39% | vRNP binding region | no |
| A/duck/Cambodia/083D1/2011 | M1 | 580 | G | T | Gly194Stop | stop gained | 2.70% | vRNP binding region | no |
| A/duck/Cambodia/083D1/2011 | M1 | 596 | C | T | Ala199Val | nonsynonymous | 2.92% | vRNP binding region | no |
| A/Cambodia/X0128304/2013 | M1 | 619 | G | A | Gln198Gln | synonymous | 5.85% | vRNP binding region | no |
| A/duck/Cambodia/PV027D1/2010 | M1 | 621 | C | T | Ala199Val | nonsynonymous | 4.56% | vRNP binding region | no |
| A/Cambodia/X0128304/2013 | M1 | 638 | A | G | Ile205Val | nonsynonymous | 2.67% | vRNP binding region | no |
| A/Cambodia/X0207301/2013 | M1 | 679 | G | A | Gly220Gly | synonymous | 6.28% | vRNP binding region | no |
| A/Cambodia/X1030304/2013 | M1 | 703 | G | A | Ala227Thr | nonsynonymous | 1.88% | vRNP binding region | no |
| A/Cambodia/X1030304/2013 | M1 | 717 | T | C | Asp231Asp | synonymous | 3.05% | vRNP binding region | no |
| A/Cambodia/X0128304/2013 | M1 | 742 | C | G | Ala239Ala | synonymous | 5.10% | vRNP binding region | no |
| A/Cambodia/X1030304/2013 | M1 | 751 | A | C | Arg243Arg | synonymous | 2.61% | vRNP binding region | no |
| A/Cambodia/X0128304/2013 | M2 | 742 | C | G | Pro10Arg | nonsynonymous | 5.10% | No known function | no |
| A/Cambodia/X1030304/2013 | M2 | 751 | A | C | Lys13Asn | nonsynonymous | 2.61% | No known function | no |
| A/Cambodia/X1030304/2013 | M2 | 861 | G | A | Cys50Tyr | nonsynonymous | 2.03% | A Cys at position 50 is a palmitoylation site that enhances virulence. | yes |
| A/Cambodia/X0125302/2013 | M2 | 882 | A | G | Arg61Gly | nonsynonymous | 2.28% | No known function | no |
| A/Cambodia/V0417301/2011 | M2 | 955 | A | C | Asp85Ala | nonsynonymous | 4.07% | No known function | no |
| A/Cambodia/V0401301/2011 | M2 | 978 | G | A | Trp89Stop | stop gained | 6.87% | No known function | no |
| A/Cambodia/X0125302/2013 | NS1 | 298 | A | G | Glu92Glu | synonymous | 2.49% | An Asp at 92 helps to confer cytokine resistance in H5N1 viruses. | yes |
| A/Cambodia/X0128304/2013 | NS1 | 302 | A | G | Glu92Glu | synonymous | 29.06% | An Asp at 92 helps to confer cytokine resistance in H5N1 viruses. | yes |
| A/Cambodia/X1030304/2013 | NS1 | 379 | A | G | Lys121Lys | synonymous | 1.93% | Part of NS1 effector domain, which interacts with many host proteins | yes |
| A/Cambodia/X0128304/2013 | NS1 | 413 | C | T | Phe129Phe | synonymous | 1.77% | Part of NS1 effector domain, which interacts with many host proteins | yes |
| A/Cambodia/X0128304/2013 | NS1 | 502 | C | T | Pro159Leu | nonsynonymous | 2.88% | Part of the NS1 nuclear export signal mask. | yes |
| A/Cambodia/X0128304/2013 | NS1 | 554 | C | T | Leu176Leu | synonymous | 3.06% | Part of NS1 effector domain, which interacts with many host proteins | yes |
| A/Cambodia/X0207301/2013 | NS1 | 609 | A | G | Arg199Arg | synonymous | 4.59% | Part of NS1 effector domain, which interacts with many host proteins | yes |
| A/duck/Cambodia/Y0224301/2014 | NS1 | 646 | T | C | Leu207Pro | nonsynonymous | 2.22% | NS1 flexible tail, which interacts with host machinery. | yes |
| A/duck/Cambodia/Y0224301/2014 | NS1 | 654 | C | T | Pro210Ser | nonsynonymous | 2.55% | NS1 flexible tail, which interacts with host machinery. | yes |
| A/Cambodia/X0128304/2013 | NEP | 554 | C | T | Ser24Leu | nonsynonymous | 3.06% | No known function. | no |
| A/Cambodia/X0207301/2013 | NEP | 609 | A | G | Glu47Gly | nonsynonymous | 4.59% | This site was implicated in enhanced virulence of H5N1 in ferrets. | yes |
| A/duck/Cambodia/Y0224301/2014 | NEP | 646 | T | C | Phe55Leu | nonsynonymous | 2.22% | No known function | no |
| A/duck/Cambodia/Y0224301/2014 | NEP | 654 | C | T | Ser57Ser | synonymous | 2.55% | No know function | no |
